# Nuclear receptor E75/NR1D2 drives tumor malignant transformation by integrating Hippo and Notch pathways

**DOI:** 10.1101/2024.06.27.600913

**Authors:** Xianping Wang, Yifan Guo, Peng Lin, Min Yu, Sha Song, Wenyan Xu, Du Kong, Yin Wang, Yanxiao Zhang, Fei Lu, Qi Xie, Xianjue Ma

**Affiliations:** Westlake Laboratory of Life Sciences and Biomedicine, Hangzhou 310024, Zhejiang, China; School of Life Sciences, Westlake University, Hangzhou 310024, Zhejiang, China; Institute of Biomedicine and Biotechnology, Shenzhen Institute of Advanced Technology, Chinese Academy of Sciences, Shenzhen, China; State Key Laboratory of Chemical Oncogenomics, Key Laboratory of Chemical Genomics, Peking University Shenzhen Graduate School, Shenzhen 518055, China

**Keywords:** tumor, steroid hormone, E75, NR1D2, Notch signaling, Hippo signaling, glioblastoma, glioblastoma stem cell, CUT&Tag, *Drosophila*

## Abstract

Hormone therapy resistance and the ensuing aggressive progression of tumors present a significant clinical challenge. However, the mechanisms underlying the induction of tumor malignancy by hormone inhibition remain poorly understood. Here we show that *Drosophila* malignant epithelial tumors exhibit a comparable decrease in ecdysone signaling, the primary steroid hormone pathway in *Drosophila*. Furthermore, we find that ectopic expression of the nuclear receptor *E75* partially mimics ecdysone signaling inhibition, specifically promoting the malignant transformation of benign tumors by integrating the Hippo and Notch signaling pathways. Genome-wide DNA binding profiles and biochemistry data reveal that E75 not only physically interacts with the transcription factors of both Hippo and Notch pathways, but also exhibits widespread co-binding to their target genes, thus contributing to tumor malignancy. Moreover, we validate these findings by demonstrating that depletion of *NR1D2*, the mammalian equivalent of E75, inhibits the activation of Hippo and Notch target genes, impeding glioblastoma progression *in vivo*. In summary, our study unveils a previously unrecognized mechanism by which hormone inhibition promotes tumor malignant transformation, while providing a conserved mechanistic understanding of how Hippo and Notch pathways are integrated by the oncogene E75/NR1D2 during tumor progression.

## Introduction

Hormones play a pivotal role in controlling cell division and growth, and their dysregulation can lead to uncontrolled cell proliferation and tumor formation.^1–3^ Many hormones can function as growth factors, thereby stimulating cell proliferation. For instance, estrogen has been shown to promote the growth of hormone receptor-positive breast cancer cells, and women who had menopausal hormone therapy shortly after menopause have been found to have a significantly increased risk of developing invasive breast cancer.^4^ Similarly, the male sex hormone androgens can stimulate prostate cancer cells to grow through the activation of androgen receptor. Consequently, hormone therapy is widely employed as a treatment strategy for both breast and prostate cancers, yielding highly effective initial responses in terms of impeding tumor growth.^1,5,6^ However, it is important to note that the development of resistance to hormone therapy poses a significant clinical problem.^1,7^ In the case of breast cancer, around 30-40% of patients experience this resistance.^8^ More strikingly, most male patients undergoing androgen deprivation therapy eventually progress to advanced castration-resistant prostate cancer (CRPC).^9^ The resistant tumor cells have increased proliferation ability and becomes more aggressive, and patients have a worse survival outcome.^10,11^ Nonetheless, the underlying mechanisms that trigger the malignant transformation in hormone signaling-reduced tumor cells remain largely unexplored.

*Drosophila melanogaster* is a widely used model organism in cancer research due to its genetic tractability and the high degree of conservation of key cancer-related pathways.^12–15^ The evolutionarily conserved Hippo and Notch signaling pathways, both initially discovered and characterized in *Drosophila*, play crucial roles in cell proliferation, differentiation, and apoptosis.^16,17^ The activation of the transcription complexes of Hippo and Notch pathways, namely the YAP/TAZ-TEAD and NICD-RBPJ-MAML complexes, respectively, promotes tumor progression and malignancy, including glioblastoma (GBM), a highly aggressive type of brain cancer.^18,19^ Previous studies have uncovered a complex interplay between these two pathways. For instance, activation of YAP/TAZ can increase the expression of Notch receptors and ligands, thereby activating Notch signaling.^20^ Conversely, Notch signaling can inhibit the Hippo pathway to increase the activity of YAP/TAZ, thus promoting cell proliferation and inhibiting apoptosis.^21^ However, the precise molecular mechanisms underlying the integration of the Hippo and Notch pathways during cancer pathogenesis remain incompletely understood.

The hormonal system of *Drosophila*, while less complex than that of mammals, exhibits several similarities to its mammalian counterparts. As a result, *Drosophila* serves as a valuable tool in the field of hormonal research.^14^ The investigation of ecdysone, a prominent steroid hormone in *Drosophila*, has shed light on its diverse roles in both physiological and pathological circumstances, ranging from the regulation of intestinal stem cell fate and morphogenesis to collective cell migration and tumor progression.^22–26^ Ecdysone interacts with and activates ecdysone receptor (EcR), which in turn binds to specific regions of the DNA known as ecdysone response elements (EcREs), thus initiating a cascade of ecdysone-responsive transcription factor expression. The early ecdysone-responsive gene *Eip75B* (*E75*) encodes a heme-binding nuclear receptor and has been implicated in various biological processes, including cell growth and differentiation,^22,23,27^ whereas the roles and mechanisms of *Eip75B* in tumorigenesis remain unknown.

In this study, we compared the transcriptome difference between benign and malignant *Drosophila* epithelium tumors and discovered that ecdysone signaling in specifically inhibited in malignant tumors. We further showed that blocking the ecdysone signaling by ectopic expression of *E75* transforms benign tumors into malignant ones. E75 drives cell proliferation and tumor malignancy by integrating Hippo and Notch signaling pathways at the transcription factor level. Moreover, we demonstrated that *NR1D2*, the mammalian ortholog of *E75*, has a functionally and mechanically conserved role in regulating GBM progression *in vivo*.

## Results

### Ecdysone signaling is inhibited in malignant tumor

*Drosophila* tumors can be classified into two main subtypes: neoplastic and hyperplastic.^15^ Neoplastic tumors exhibit signs of cell polarity loss and differentiation defect, resulting the formation of multi-layered invasive tumors. The typical genes of this category include *scribble* (*scrib*), *discs large* (*dlg*), and *lethal giant larvae* (*lgl*).^15,28^ Conversely, hyperplastic tumors maintain the cell architecture of the single-layered epithelium and preserve the cell fate. Examples of these tumor regulating genes include *warts/lats* (*wts*) and oncogenic *Ras* (*Ras^V^*^12^). We generated GFP labeled clones harboring mutations of different tumor suppressor genes and oncogenes using the mosaic analysis with a repressible cell marker (MARCM) system (Fig. 1A).^29,30^ In the presence of surrounding wild type (WT) cells, the *scrib* mutant clones undergo elimination through a conserved process known as tumor suppressive cell competition (Figs. 1A and 1B).^29,31–34^ However, when *Ras^V12^* is co-expressed, the *scrib* clones are transformed into malignant tumors (Figs. 1B and 1B’).^29,35,36^ Interestingly, while mutation of *wts* alone resulted in larger clones compared to *Ras^V12^* overexpression (Fig. 1B), co-deletion of *wts* failed to transform *scrib* clones into large tumor masses (Figs. 1B and 1B’).

**Fig. 1.**
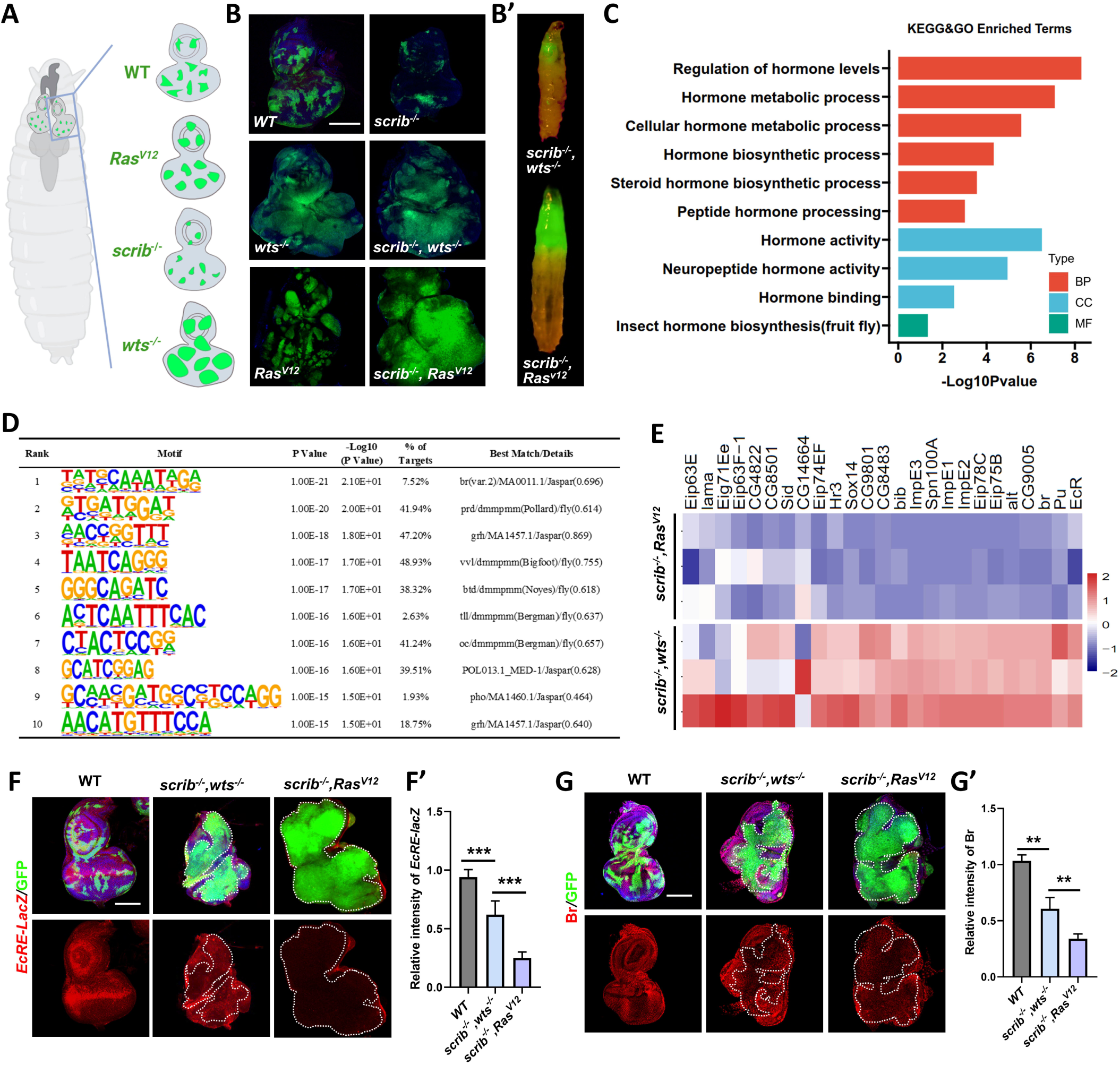
Ecdysone signaling is inhibited in malignant tumor. (**A**) Schematic representation of eye-antennal discs from a *Drosophila* larva of *ey-Flp-*MARCM-induced mosaics. (**B**) Eye-antennal discs bearing *ey-Flp-*MARCM-induced mosaics of indicated genotypes. (B’) Representative images of tumor-bearing larvae of *scrib^−/−^, wts^−/−^* and *scrib^−/−^, Ras^V12^.* (**C**) Enrichment of hormone related terms between *scrib^−/−^, wts^−/−^* and *scrib^−/−^, Ras^V12^* tumors from KEGG and GO analyses. (**D**) HOMER motif analysis of ecdysone-related genes in *scrib^−/−^, wts^−/−^* and *scrib^−/−^, Ras^V12^* tumors. (**E**) Heatmap profiles of ecdysone signaling-related genes in *scrib^−/−^, wts^−/−^* and *scrib^−/−^, Ras^V12^* tumors. (**F-G**) Confocal images of eye-antennal discs bearing *ey-Flp-*MARCM-induced mosaics of indicated genotype were stained with anti-β-galactosidase antibody for the *EcRE-LacZ* staining (F) or anti-Br antibody (G). Quantification of relative intensity of *EcRE-LacZ* (F’, from left to right, n = 9, 8, 8) and Br (G’, from left to right, n = 5, 8, 11) in GFP positive mosaics clones. Statistical analysis by Ordinary one-way ANOVA test; mean ± SD. \*\*\*\**p* < 0.0001. Scale bars: 100μm (B, F, G).

To investigate the mechanisms underlying the phenotypic differences between the *scrib^−/−^, wts^−/−^* and *scrib^−/−^, Ras^V12^* tumors (Fig. 1B’), we conducted bulk RNA-seq analysis. Interestingly, by analyzing the differentially expressed genes (DEGs), we noticed significant enrichment of multiple hormone related terms (Figs. 1C and S1A). We further performed motif enrichment analysis using HOMER (Hypergeometric Optimization of Motif EnRichment)^37^ to identify the crucial transcription factors (TFs) involved in the regulation of the DEGs. Of note, the highest-ranked TF identified was *broad* (*br*) (Fig. 1D), a known “early” ecdysone induced gene that subsequently triggers the activation of “late” genes in the ecdysone signaling. Moreover, the Gene Set Enrichment Analysis (GSEA) and RNA-seq analysis both demonstrated a significant down-regulation of the ecdysone signaling in the *scrib^−/−^, Ras^V12^* malignant tumor compared to the *scrib^−/−^, wts^−/−^* benign tumors (Figs. 1E and S1B). Consistent with these findings, the transcriptional activation of ecdysone signaling *in vivo* was severely impeded in malignant tumors, but only mildly inhibited in benign tumors, as demonstrated by the ecdysone response element (*EcRE*)-driven LacZ (*EcRE-LacZ*) reporter and Br staining (Figs. 1F-G’). Another malignant tumor induced by *lgl^−/−^, Ras^V12^* also exhibited a significant reduction in Br (Figs. S1C and S1C’). In line with the potential tumor-suppressive role of increased ecdysone signaling, we observed a robust suppression of *lgl^−/−^, Ras^V12^*-induced tumorigenesis and restoration of the pupation defect upon hyperactivation of ecdysone signaling through ectopic expression of two different isoforms of *EcR* (*EcRA* and *EcRB1*) (Figs. S1D-D”). Collectively, these findings indicate that the inhibition of ecdysone signaling plays a critical role in the malignant transformation of tumors.

### E75 overexpression induces tumor malignant transformation

One of the extensively studied early ecdysone-induced genes, E75, has been genetically implicated as a repressor of the ecdysone-triggered cascade.^38,39^ In line with this, *ey-Flp*- MARCM-induced clonal overexpression of the A isoform of *Eip75B* (abbreviated as *E75*) significantly inhibited the activation of endogenous ecdysone signaling (Figs. 2A-B). Differentiation failure is one of the key characteristics that distinguish malignant from benign tumors. Clones overexpressing *E75* exhibited a dramatic defect in differentiation, as evidenced by the staining of the neuronal differentiation marker Elav (Fig. 2C). Consistent with its potential role in promoting malignancy, we found that the overexpression of *E75* exhibited a synergistic effect when combined with either *scrib^−/−^* or *wts^−/−^*, resulting in tumor overgrowth and pupation defects (Figs. 2D-D”). Intriguingly, the overexpression of *E75* not only significantly increased the size of *scrib^−/−^, wts^−/−^*-induced benign tumors (Figs. 2D and 2D’), but also astonishingly transformed them into malignant tumors, resulting in 96.3% of the animals being unable to pupate and instead developing into giant larvae (Figs. 2D and 2D’’). This transformation was accompanied by a significant enhancement in tumor invasive ability, as evidenced by the abundant Mmp1 staining observed at the leading edge of the invasive tumor cells (Figs. 2E and 2E’). Moreover, *E75* overexpression also synergistically transformed other benign tumors into malignant ones, including *Ras^V12^*and *Raf^GOF^, scrib^−/−^* (Fig S2). Taken together, these results suggest that ectopic expression of *E75* could facilitate the tumor malignant transformation.

**Fig. 2.**
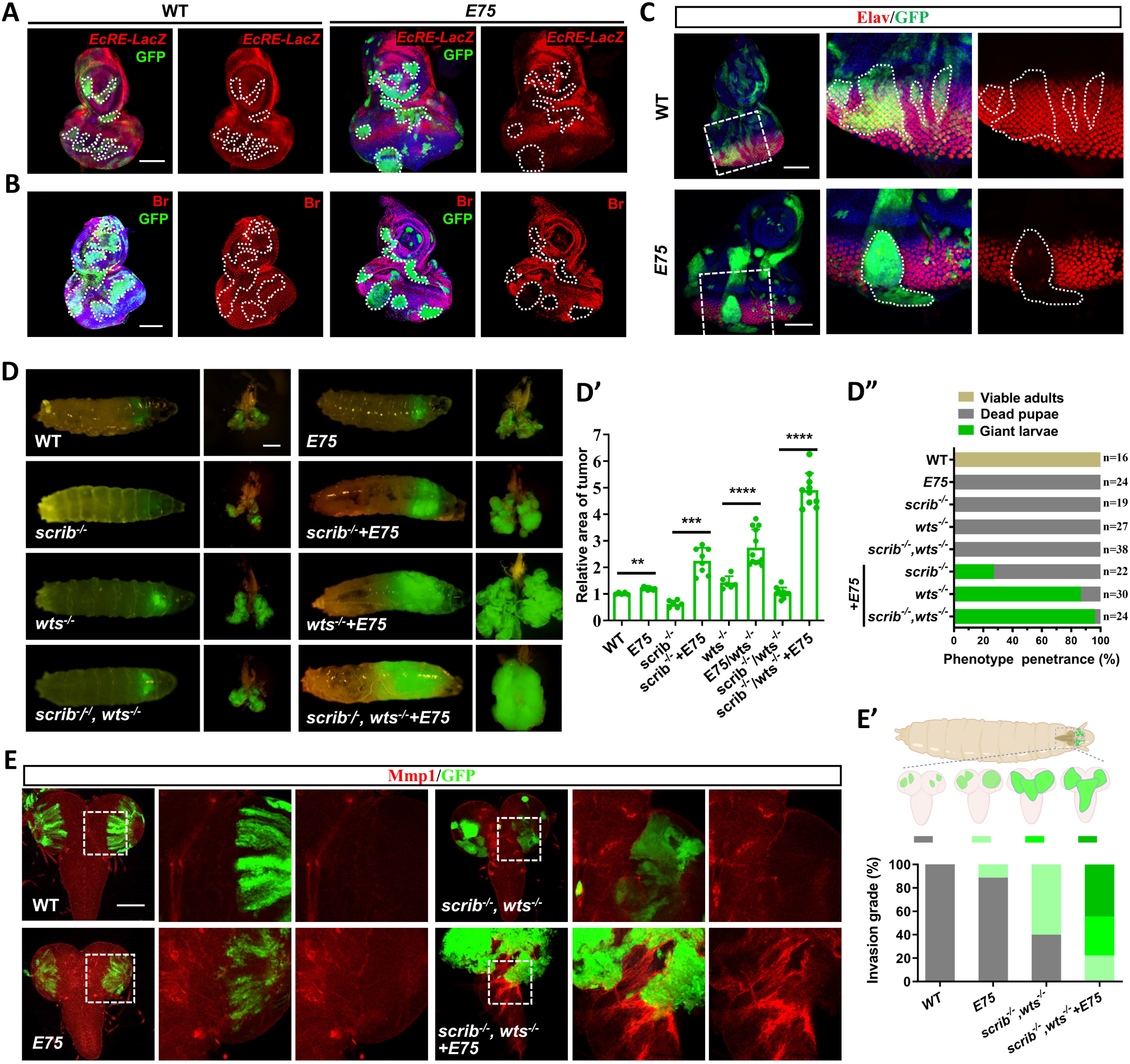
E75 overexpression induces tumor malignant transformation. (**A-C**) Confocal images of eye-antennal discs bearing *ey-Flp-*MARCM-induced mosaics of wild-type and E75 overexpression were stained with anti-β-galactosidase antibody for the *EcRE-LacZ* staining (A), anti-Br antibody (B), or anti-Elav antibody (C). Clones are circled by the white dash line. (**D**) Dorsal views of *ey-Flp*-MARCM-induced GFP-positive tumor-bearing larvae and the corresponding eye disc or tumor (right). Quantification of relative tumor size (D’) (*n* = 5, 10, 6, 8, 6, 12, 10, 10, from left to right) and larvae pupation rate (D”). (**E**) Representative confocal images of the ventral nerve cord (VNC) from *ey-Flp*-MARCM- induced tumors with indicated genotype stained with anti-Mmp1 antibody. Quantification of the invasion grade of each genotype (E’, bottom panel). Carton illustration of the invasion grade of VNC in tumor-bearing larvae (E’, top panel) (from left to right, n=11,9, 10, 9). Statistical analysis by Ordinary one-way ANOVA test; mean ± SD. \*\*\*\**p* < 0.0001. Scale bars: 100 μm (A, B, C, E), 200 μm (D).

### E75 inactivates the Hippo pathway to induce tumor malignancy

To further explore the underlying mechanisms by which *E75* overexpression promotes tumor malignancy, we performed bulk RNA-seq analysis on tumors derived from *scrib^−/−^, wts^−/−^* and *scrib^−/^, wts^−/−^+E75*. A total of 3457 DEGs were identified, including 1985 upregulated genes and 1472 downregulated genes (Fig. S3A). We subjected these DEGs to further analysis using the Kyoto Encyclopedia of Genes and Genomes (KEGG) and observed a significant enrichment and inactivation of the Hippo signaling pathway (Figs. 3A, S3B, and S3C), which is a conserved pathway crucial for size control and tumorigenesis.^16^ The overexpression of *E75* resulted in an increase in clone size (Figs. 3B and 3B’). Conversely, the depletion of *E75* not only decreased clone size, adult wing, and eye size under physiological conditions, but also inhibited tumor overgrowth induced by *scrib^−/^, wts^−/−^* (Figs. 3B, 3B’, and S3D). Consistently, E75 overexpression or depletion could up-regulate or downregulate the expression of multiple target genes in the Hippo pathway, including Wingless (Wg), *expanded* (*ex*), Cyclin E (CycE), and *bantam* (*ban*) (Fig. 3C). Together, these data indicate that E75 negatively regulates Hippo signaling.

**Fig. 3.**
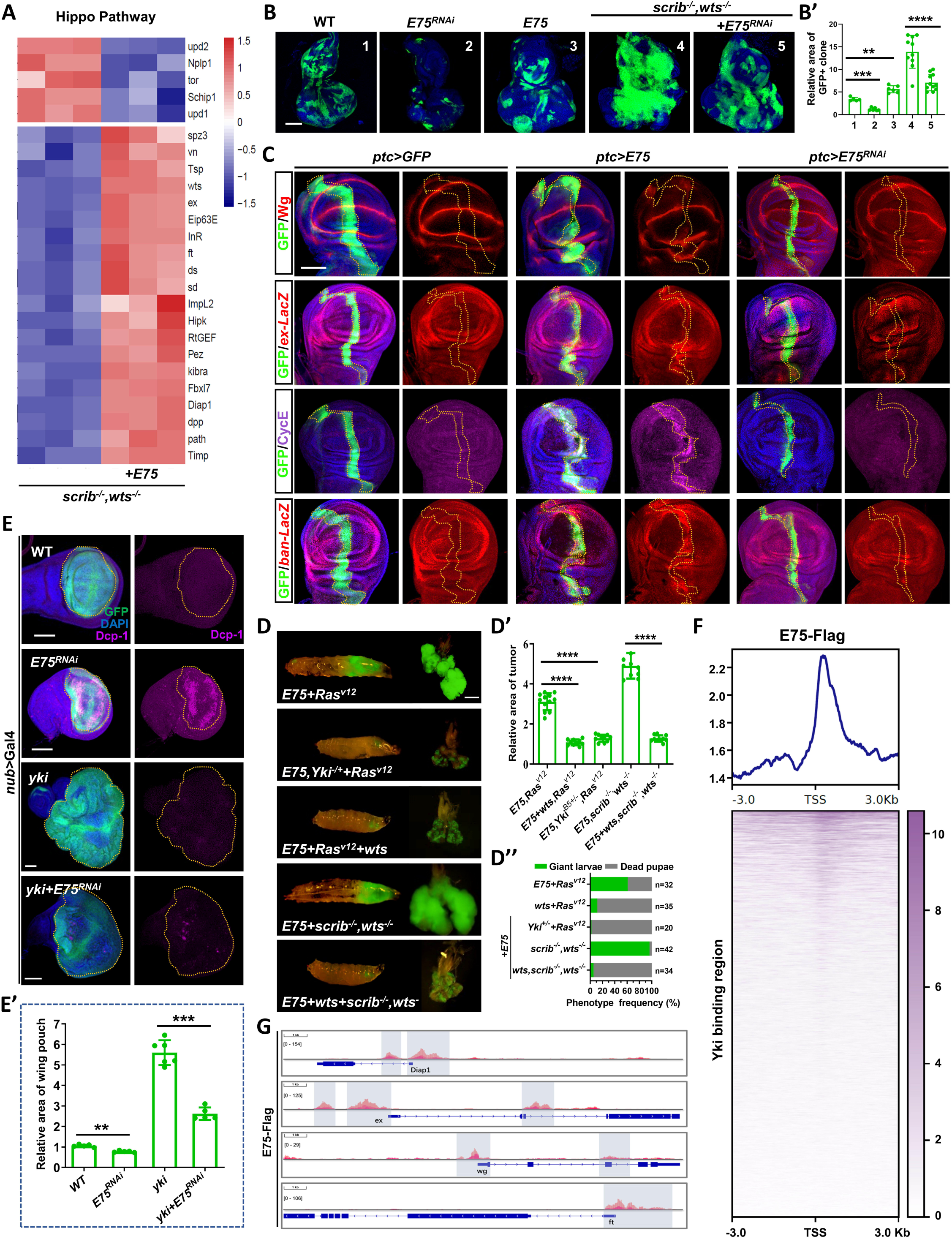
E75 inactivates the Hippo pathway to induce tumor malignancy. (**A**) Heatmap profiles of the expression of Hippo signaling-related genes in *scrib^−/−^, wts^−/−^* and *E75, scrib^−/−^, wts^−/−^* tumors. (**B**) Representative images of eye-antennal discs bearing *ey-Flp-*MARCM-induced mosaics of indicated genotypes. (B’) Quantification of the relative size of GFP positive regions in (B) (*n* = 5, 6, 6, 10, 11). (**C**) Confocal images of the wing disc with E75 overexpression or knockdown under the control of *ptc* promoter were stained with anti-Wg, anti-CycE, or anti-β-galactosidase antibody for *ex-LacZ* and *ban-LacZ* staining. (**D**) Dorsal views of *ey-Flp*-MARCM-induced GFP-positive tumor-bearing larvae and the corresponding tumor. Quantification of relative tumor size (D’) (n = 12, 10, 10, 10, 10) and larvae pupation rate (D”). (**E**) Confocal images of wing disc with indicated genotypes were stained with anti-Dcp-1 antibody. (E’) Quantification of the relative size of wing pouch region of D (n=5, 5, 6, 5). (**F**) Line plots of the average Cut&Tag signal of E75 peaks (top panel) and heatmaps of the Cut&Tag signals of Yki in *Drosophila* (bottom panel). Cut&Tag signals are displayed within a region spanning ±3 kb around all canonical transcription start sites (TSS) genome-wide. (**G**) Browser shots of E75 CUT&Tag signal at canonical Hippo pathway target genes. The shaded areas correspond to E75 peaks. Statistical analysis by Ordinary one-way ANOVA test (D’) or Brown-Forsythe and Welch ANOVA tests (B’, E’); \*\**p* < 0.01, *** *p* < 0.001, **** *p* < 0.0001; Scale bars: 100 μm (B, C, E) and 200um (D).

In *Drosophila*, the core components of the Hippo pathway consist of Hippo (Hpo), Warts (Wts), and Yorkie (Yki). Hpo phosphorylates Wts, which subsequently phosphorylates and inactivates the transcriptional coactivator Yki.^16^ Consistent with the observed enrichment of Hippo signaling in E75-induced malignant tumors, the Hippo pathway was also found to be enriched through a bulk RNA-seq analysis performed on *E75*-overexpressed wing discs. (Figs. S3E and S3F). To investigate the relationship between E75 and the Hippo pathway components, we conducted genetic epistasis analysis. Overexpression of *E75*, driven by the *patched* (*ptc*) promoter, efficiently induced overgrowth and upregulated CycE. These effects were significantly suppressed upon knockdown of *yki* or coexpression of *wts* (Figs. S3G and S3G’). Similarly, reducing *yki* activity significantly blocked the tumor malignancy and pupation defect induced by E75 overexpression in both *Ras^V12^* and *scrib^−/^, wts^−/−^* under pathological conditions (Figs. 3D-D”). On the other hand, when *E75* was depleted, the ectopic expression of *yki*-induced overgrowth phenotype was also impeded (Figs. 3E and 3E’). Hence, these findings collectively suggest that E75 acts genetically in parallel with Yki.

Given that *E75* possesses a conserved DNA binding domain (DBD) and the ability to initiate gene transcription, we subsequently performed Cleavage Under Targets and Tagmentation (CUT&Tag) analysis^40^ to investigate the genome-wide DNA binding profiles of E75 in dissected wing pouch regions from *E75* overexpressed discs (Fig. S3H). In addition to the well-established binding motif, we also identified novel binding motifs of *E75* using the HOMER software (Fig. S3I). By integrating the public available chromatin immunoprecipitation sequencing (ChIP-seq) data of Yki-bound chromatin, ^41^we found that the CUT&Tag signal of E75 is significantly enriched in the region where Yki is bound (Fig. 3F). Supporting this, our analysis revealed numerous E75 binding sites located within the promoter regions of canonical Hippo target genes, such as *Death-associated inhibitor of apoptosis 1* (*Diap1*), *ex*, *wg*, and *fat* (*ft*) (Fig. 3G). These findings suggest that E75 may serve as a potential partner of Yki in the transcriptional regulation of downstream genes.

### E75 activates the Notch pathway to promote tumor malignancy

It is noteworthy that the activation of Yki alone cannot comprehensively explain the malignant transformation of *scrib^−/−^, wts^−/−^*benign tumors. This is because the deletion of *wts* alone is sufficient to hyperactivate Yki, indicating the involvement of additional signaling pathways essential for E75-induced malignant transformation. Of note, in addition to *Ras^V12^*, *scrib*-mutated cells could also synergize with the oncogenic form of *Notch* to induce a massive overgrowth.^29^ Interestingly, upon re-analysis of RNA-seq data obtained from *scrib^−/^, wts^−/−^+E75* tumors, a significant enrichment of the Notch signaling was observed (Figs. S4A and S4B). Furthermore, the examination of the binding regions of *Suppressor of Hairless* [*Su(H)*], the transcription factor of Notch pathway, revealed a strong CUT&Tag signal of E75 (Fig. S4C). Additionally, several genes known to be targeted by Notch, such as *E(spl)m*β*- HLH*, *E(spl)m*α*-BFM*, *E(spl)m7-HLH*, and *E(spl)m8-HLH*, displayed binding sites for E75 (Fig. 4A). Consistent with these findings, the ectopic expression of *E75* upregulated the expression of several Notch target genes in both physiological and E75-induced tumorigenic conditions (Figs. 4B, 4C, S4D, and S4E), while the depletion of *E75* inhibited their endogenous expression (Figs. 4B and S4D).

**Fig. 4.**
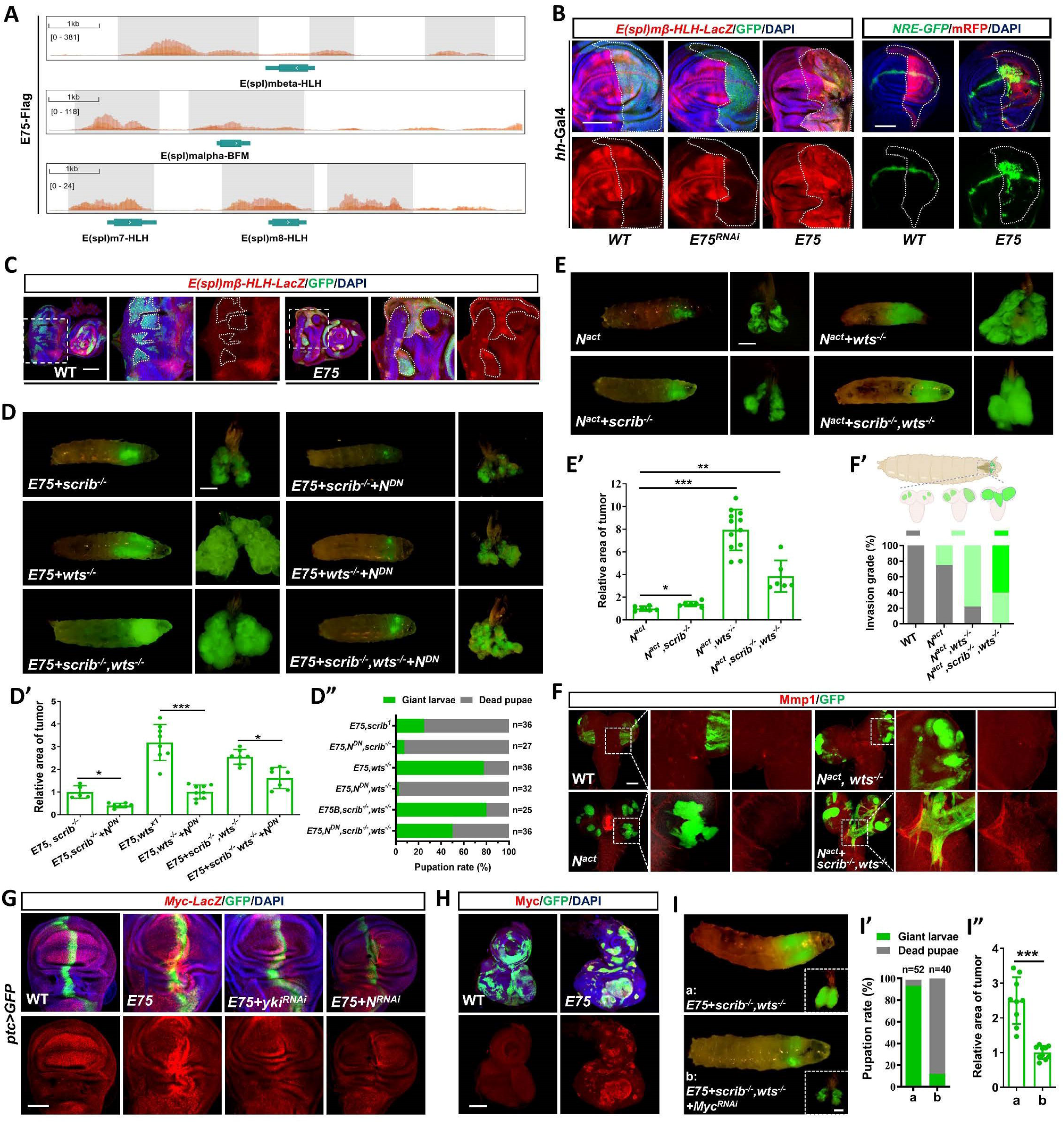
E75 activates the Notch pathway to promote tumor malignancy. (**A**) Browser shots of E75 CUT&Tag signal at canonical Notch pathway target genes. The shaded areas correspond to E75 peaks. (**B**) Confocal images of wing disc with indicated genotypes were stained with anti-β- galactosidase antibody for *E(spl)m*β*-HLH-LacZ* staining (left) or *NRE-GFP* expression (right). (**C**) Eye-antennal discs bearing *ey-Flp-*MARCM-induced mosaics of wild-type and E75 overexpression were stained with anti-β-galactosidase antibody for the *E(spl)m*β*-HLH-LacZ* staining. (**D, E**) Dorsal views of *ey-Flp*-MARCM-induced GFP-positive tumor-bearing larvae and the corresponding eye disc or tumor (right). Quantification of relative tumor size (D’, *n* = 12, 13, 13, 14, 10, 10), (E’, n = 8, 10, 16, 12), and larvae pupation rate (D”). (**F**) Representative confocal images of VNC from *ey-Flp*-MARCM-induced tumors with indicated genotype were stained with anti-Mmp1 antibody. Carton illustration of invasion grade of VNC in tumor-bearing larvae (F’, top panel). Quantification of invasion percentage of each genotype (F’, bottom panel) (n = 12, 8, 10, 9). (**G**) Representative confocal images of wing disc bearing indicated genotypes were stained with anti-β-galactosidase antibody for *Myc-LacZ* staining. (**H**) Eye-antennal discs bearing *ey-Flp-*MARCM-induced mosaics of wild-type and E75 overexpression were stained with anti-Myc antibody. (**I**) Dorsal views of *ey-Flp*-MARCM-induced GFP-positive tumor-bearing larvae and the corresponding eye disc or tumor (right corner). Quantification of larvae pupation rate (I’) and relative tumor size (I’’) (*n* = 10, 12). Statistical analysis by Student’s t-test (I”) or Brown-Forsythe and Welch ANOVA tests (D’, E’); \**p* < 0.05, **** *p* < 0.0001; Scale bars: 100 μm (B, C, F, G, H) and 200um (D, E, I).

To further verify the *in vivo* role of Notch in E75-induced tumor malignancy, we inhibited Notch activation by expressing a dominant negative form of *Notch* (*N^DN^*). Remarkably, Notch inhibition dramatically suppressed the synergistic tumor promoting effect caused by E75 overexpression and restored the pupation defects (Figs. 4D-D”). On the contrary, genetically activating the Notch pathway by co-expressing the activated form of the Notch intracellular domain (*N^act^*) could transform *scrib^−/^, wts^−/−^* benign tumors into malignant ones and lead to the development of aggressive tumors (Figs. 4E-F’). It is noteworthy that the size of *N^act^+wts^−/−^* tumor looks generally larger than that of *N^act^+scrib^−/−^, wts^−/−^* tumor, probably due to the two-dimensional expansion of tumor cells without disrupting the single-layer polarity architecture of the eye epithelium. These findings collectively suggest that the activation of the Notch pathway is both necessary and sufficient for the malignant transformation of *E75*-mediated tumor progression.

### Myc is an essential downstream gene for E75-induced tumor malignancy

The *Myc* oncogene plays a critical role in the progression of various human cancers. Intriguingly, *Myc* also serves as an evolutionally conserved transcriptional target for both the Hippo and Notch signaling pathways.^42,43^ Given this, we further explored whether Myc acting as a crucial downstream target gene in the context of tumor malignancy induced by E75 overexpression. Depletion of *E75* decreased endogenous *Myc* transcription (Fig. S4F), whereas ectopic expression of *E75* caused a robust increase in both transcription and protein levels of *Myc*, in a Yki- and Notch-dependent manner (Figs. 4G and 4H). In line with this, our CUT&Tag analysis identified multiple E75 binding sites within the promoter and gene body regions of *Myc* (Fig. S4G). Moreover, the depletion of *Myc* not only suppressed the tissue overgrowth phenotype caused by E75 overexpression (Figs. S4H-I’), but also significantly blocked E75-induced malignant transformation of *scrib^−/−^, wts^−/−^*tumors (Figs. 4I-I”). These data collectively demonstrate that *Myc* functions as a crucial downstream target gene of E75 in the regulation of both growth and malignancy.

### E75 integrates the Hippo and Notch pathways at the transcriptional factor level

The observed similarity in binding sites between E75 and Yki or Su(H), as demonstrated by our E75 CUT&Tag analysis, indicates a potential physical interaction between E75 and the transcriptional complexes associated with the Hippo and Notch pathways. To test this, we performed proximity ligation assay (PLA) in *Drosophila* wing and eye epithelium to detect the protein-protein interactions at subcellular level *in situ*.^44^ Compared to the negative controls, we observed robust positive PLA signals between Flag-tagged E75 and Myc-tagged Yki or HA-tagged Scalloped (Sd), the transcriptional factor of Hippo pathway (Figs. 5A and 5B). Similarly, PLA signals were detected between E75 and Sd in the eye-disc clones that co-expressing *E75^Flag^*, *yki*, and *Sd^HA^*(Fig. S5A). Furthermore, strong PLA signals were also observed between ectopically expressed E75 and endogenous NICD and Su(H) (Fig. 5C). As a second validation experiment, we performed co-immunoprecipitation (co-IP) assay to examine the physical interactions between E75 and above-mentioned TFs. Consistently, robust physical interactions were detected between HA-tagged Sd and Flag-tagged E75, as well as Myc-tagged E75 and Flag-tagged Su(H), in *Drosophila* S2 cells (Figs. 5D and 5E). Moreover, when ectopically expressed with Yki and NICD in the developing eye epithelium, Flag-tagged E75 demonstrated strong physical interactions with both (Fig. 5F). Together, these data suggest that E75 can physically bind with the downstream transcriptional complexes of both Hippo and Notch pathways.

**Fig. 5.**
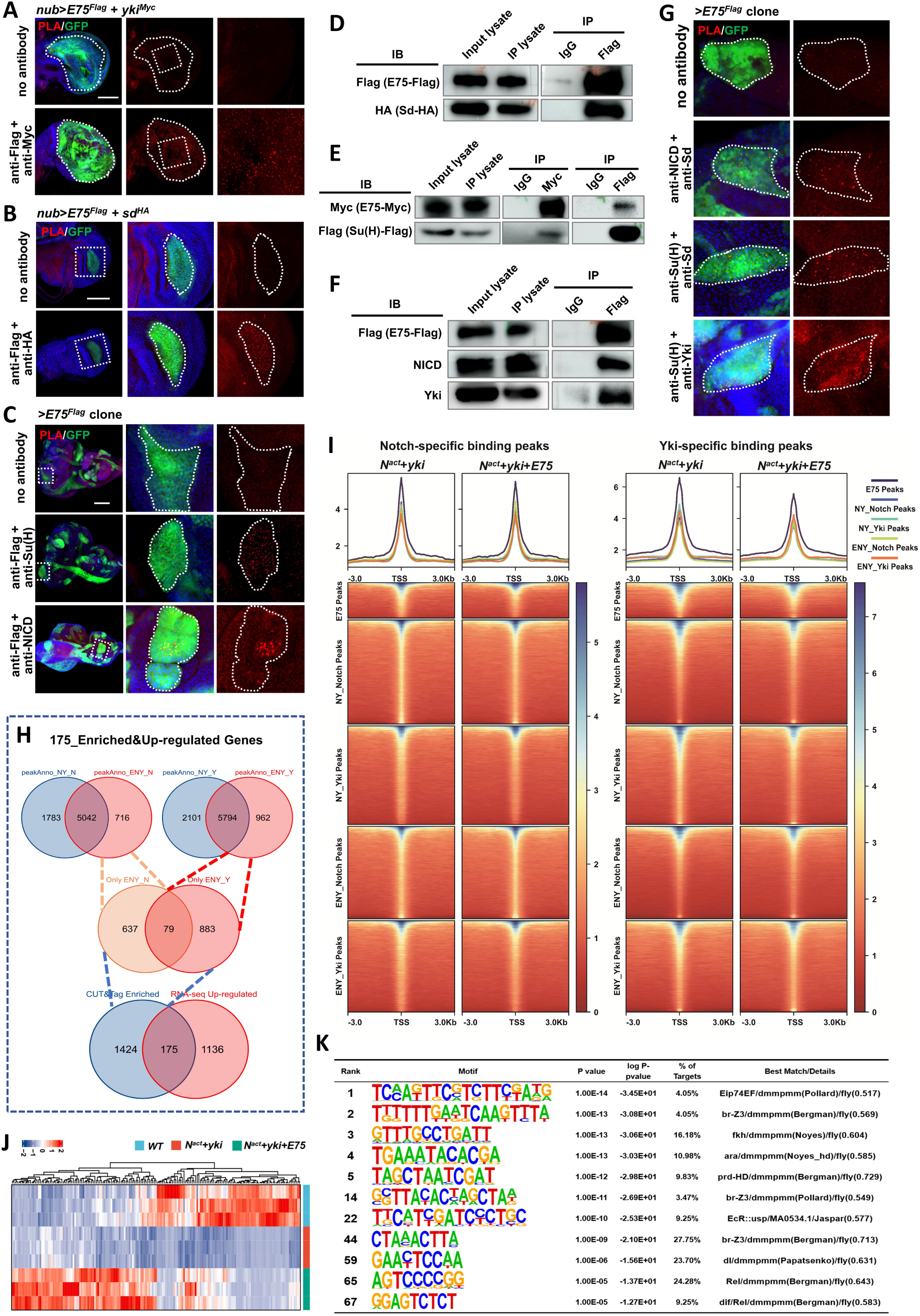
E75 integrates the Hippo and Notch pathways at the transcription factor level. (**A-B**) Proximity Ligation Assay (PLA) was performed on wing discs with indicated genotypes to test close-proximity interactions between Flag-tagged E75 and Myc-tagged Yki (A) or Flag-tagged E75 and HA-tagged Sd (B). (**C**) PLA was performed on *ey-Flp*-MARCM-induced GFP-positive clones with indicated genotypes to test close-proximity interactions between E75 and Su(H) or NICD. (**D-F**) Co-IP assay to detect physical interactions between *E75^Flag^*and *Sd^HA^*, NICD, Yki, and Su(H). Lysates from S2 cells expressing indicated constructs (D and E) or from *ey-Flp-* MARCM-induced tumors with indicated genotypes (F) were immunoprecipitated and probed with the indicated antibodies. (**G**) PLA was performed on *ey-Flp-*MARCM-induced *E75^Flag^*overexpression clones to test close-proximity interactions between endogenous NICD and Sd, Su(H) and Sd, Su(H) and Yki. (**H**) The strategy of identifying specific gene cluster that are specifically regulated by *E75* between NY (co-expression of *N^act^* and *Yki*) and ENY (co-expression of *N^act^*, *Yki,* and *E75*) groups. (**I**) Line plots (top panels) show the average Cut&Tag signal of Notch-specific peaks of *N^act^+Yki* and *E75+N^act^+Yki*, and Yki-specific peaks of *N^act^+Yki* and *E75+N^act^+Yki*. Clustered heatmaps (bottom panels) show the Notch-specific peaks of *N^act^+Yki* and *E75+N^act^+Yki* (left), and Yki-specific peaks of *N^act^+Yki* and *E75+N^act^+Yki* (right). Cut&Tag signals are displayed within a region spanning ±3 kb around all canonical TSS. E for E75, N for Notch, Y for Yki. (**J**) Heatmaps showing the relative expression level of those enriched 175 genes from H in discs with different genotypes. (**K**) HOMER motif distribution analysis of 175 upregulated genes that are enriched with peaks. Scale bars: 20um (G) and 100 μm (A, B, C).

Under tumorigenic stress, NICD can form a complex with Transcriptional coactivator with PDZ-binding motif (TAZ), the human ortholog of Yki.^21^ Indeed, we noticed that discs expressing both *yki* and *N^act^*showed a detectable but moderate PLA signal (Fig. S5B). Interestingly, it is noteworthy that no PLA singles were detected between Su(H) and Yki or Sd under physiological conditions (Fig. S5C). However, upon the overexpression of *E75*, a dramatic increase in PLA signals was observed between Su(H) and Yki or Sd, as well as NICD and Sd (Fig. 5G). This observation suggests that the forced expression of *E75* can enhance the physical association between downstream TFs of the Hippo and Notch pathways. Consistent with this notion, ectopic expression of *E75* synergistically enhanced the tumorigenic potential of the benign tumor induced by the co-expression of *N^act^*and *yki*, leading to their transformation into malignant tumors (Fig. S5D).

To further investigate the effect of E75 on the TF interacting network and chromatin landscape, we conducted NICD and Yki CUT&Tag analyses on tumor samples from *N^act^*+*yki* expressing (NY) and *E75+N^act^+yki* expressing (ENY) eye epithelium. The peaks were annotated based on to the genomic locations (Fig. S5E) and relative distance to transcriptional start sites (TSSs) (Fig. S5F). For the NY tumors, a total of 6,825-NICD annotated and 7,895 Yki-annotated peaks were identified, while for the NYE tumors, 5,758-NICD-annotated and 6,756 Yki-annotated peaks were uncovered (Fig. 5H). To visualize the levels and distribution of NICD and Yki, we generated spike-normalized coverage heatmaps at a genome-wide scale (Fig. S5G). Interestingly, we observed a general decrease in signal intensities of both NICD and Yki around the TSSs when *E75* was co-expressed (Fig. S5H). Next, we plotted the NICD and Yki-ATAC signals around the TSS of protein-coding genes where E75 bound. We found that while both NICD and Yki bind to the regions accessible to E75, their signal intensities were also reduced upon *E75* expression (Fig. 5I). The inconsistency between decreased TFs binding and increased tumor overgrowth raises the possibility that NICD and Yki may bind to additional DNA regions when E75 abundance arises.

To test this hypothesis, we conducted a comparative analysis of the NICD and Yki CUT&Tag signals in NY and ENY tumors. Our analysis revealed a total of 1,599 specific peaks within ENY tumors that were exclusively associated with either NICD or Yki (Fig. 5H). Subsequently, we performed RNA-seq analysis on tumors from NY and ENY and intersected the upregulated 1,311 genes with the 1,599 identified peaks, resulting in the identification of 175 genes (Figs. 5H and 5J). These 175 genes were then subjected to over-representation analysis (ORA), while their corresponding 300 peaks underwent transcription factor motif enrichment using the HOMER and the Find Individual Motif Occurrences (FIMO) algorithms. The HOMER enrichment analysis consistently uncovered TF binding motifs of ecdysone response-related elements (Fig. 5K), and the FIMO results revealed that the *de novo* identified E75 binding motif had the potential to bind to a range of 12.33% to 25.00% of the 300 peaks (Fig. S5I). These findings indicate that E75 and ecdysone signaling are likely to regulate these 175 genes. Intriguingly, our HOMER analysis also revealed a significant enrichment of TFs related to the Toll and Imd signaling pathways, including dorsal (dl), Relish (Rel), and Dorsal-related immunity factor (Dif) (Fig. 5K). Additionally, our KEGG analysis identified the Toll and Imd signaling pathway as one of the top significantly enriched regulons upon E75 expression (Figs. S5J and S3E). Interestingly, an increase in NICD and Yki signals was observed on the *Dif* promoter in ENY tumors (Fig. S5K), implying that *Dif* may potentially serve as an essential downstream gene responsible for the malignant transformation of ENY tumors.

Taken together, these results suggest that E75 effectively integrates the Hippo and Notch pathways at the transcriptional factor level. Ectopically expressed E75 not only physically associates with the TFs of both pathways, but also facilitates the binding of NICD and Yki to novel chromatin regions. Both actions collectively contribute to the tumor malignancy.

### Silencing *NR1D2* suppressed glioblastoma stem cell-driven tumor growth

Next, we investigated the potential role of E75 in regulating Notch and Hippo pathway-mediated tumorigenesis in mammals, with a specific focus on glioblastoma (GBM) - a highly aggressive and fatal brain tumor. We have previously demonstrated that depleting *NR1D2* (*nuclear receptor subfamily 1 group D member 2*), the mammalian ortholog of E75, impedes the mobility and viability of GBM cell lines.^45^ Consistent with our *Drosophila* tumor data, the KEGG analysis revealed significant enrichment of both the Notch and Hippo signaling pathways in *NR1D2*-depleted GBM cells (Fig. S6A).^45^ Moreover, analysis utilizing GEPIA (Gene Expression Profiling Interactive Analysis) indicated that several target genes of both pathways showed a positive correlation with elevated *NR1D2* expression in GBM (Fig. S6B). Glioblastoma stem cells (GSCs) constitute a subset of cells within the GBM that possess stem cell-like properties, they can self-renew, undergo differentiation, and contribute to therapeutic resistance.^46,47^ Notably, we found the expression level of *NR1D2* was significantly upregulated in GSCs compared to their associated neural stem cells (NSCs) (Fig. 6A).^48^ Furthermore, the depletion of *NR1D2* using two nonoverlapping short hairpin RNAs (shRNA) (Fig. S6C) strongly decreased cell proliferation of two patient-derived GSCs (MGG6 and MGG4) (Figs. 6B and S6D) and led to a significant decrease in the expression levels of Notch and Hippo pathway target genes in GSCs (Figs. 6C and 6D). Similar to the interacting network of TFs identified in *Drosophila*, we found that NR1D2 also forms physical interactions with TFs belonging to both the Hippo and Notch pathways in both GSCs and GBM cells, as demonstrated by PLA and co-IP assays (Figs. 6E-G, S6E, and S6E’). To verify whether NR1D2 could directly bind to the promoter regions of Hippo and Notch target genes, we performed chromatin immunoprecipitation (ChIP) assays in GSCs and GBM cells that stably expressed HA-tagged NR1D2 (HA-NR1D2). Our data showed that NR1D2 binds to the promoter regions of multiple Hippo and Notch target genes, including *Axl*, *HEY1*, and *TCF7* (Figs. 6H and S6F). Finally, we tested the impact of disrupting *NR1D2* on GSC-driven tumor growth *in vivo*. Luciferase-expressing GSCs, transfected with lentivirus expressing *NR1D2* shRNA or non-targeting control shRNA (shNT), were injected into the right cerebral cortex of NSG mouse brains (Fig. 6I). Bioluminescent imaging showed that *NR1D2* knockdown markedly suppressed GSC-driven tumor growth and extended the survival of mice compared with controls (Figs. 6J-K’ and S6G-H). Collectively, these results indicate that NR1D2 plays a conserved role in regulating the Hippo and Notch pathways and is crucial for GSC-induced tumor growth.

**Fig. 6.**
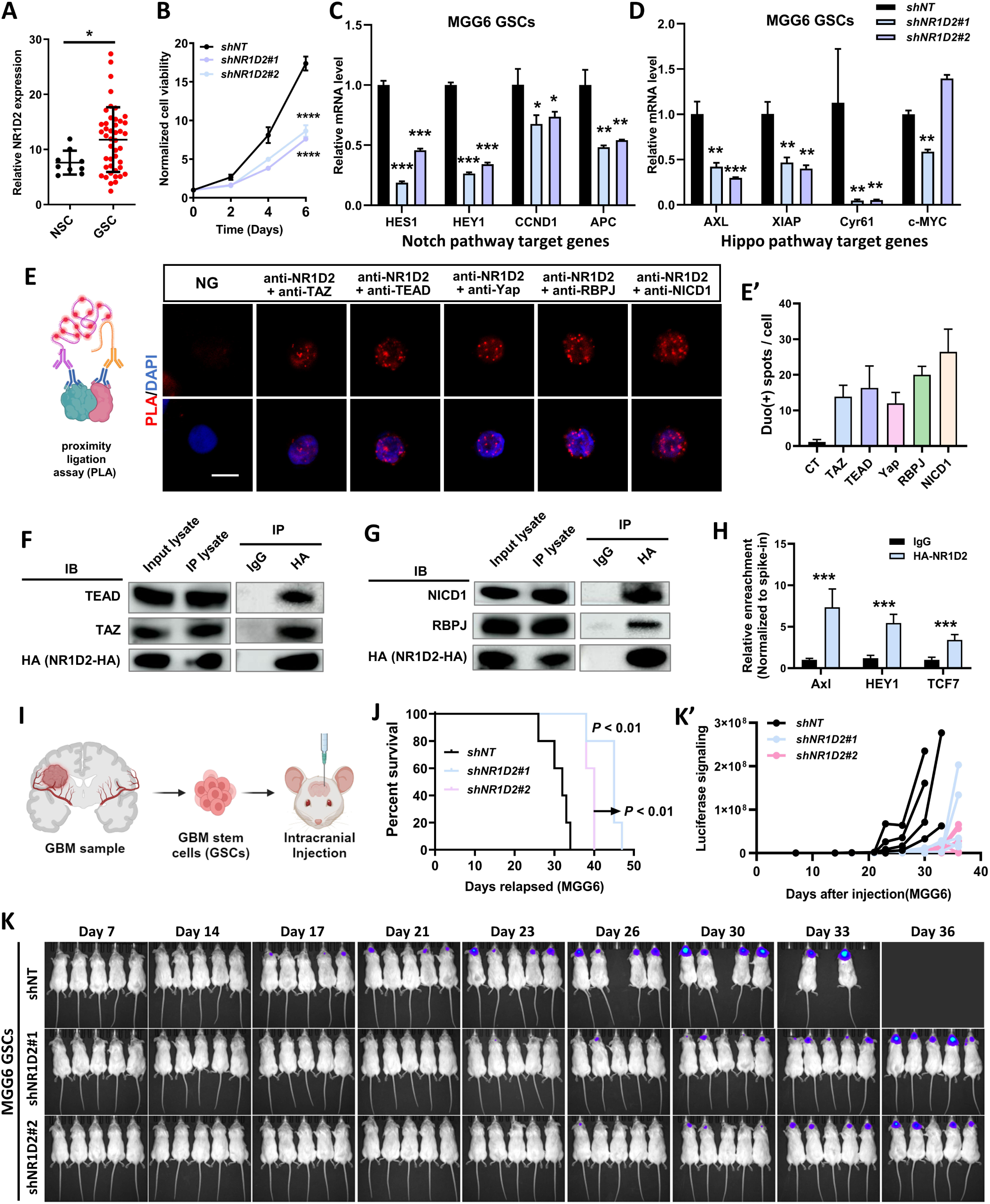
Silencing *NR1D2* suppressed glioblastoma stem cell-driven tumor growth. (**A**) Relative expression of *NR1D2* in neural stem cell (NSC) and glioblastoma stem cell (GSC). (**B**) Relative cell viability of MGG6 of shNT and *shNR1D2*. (**C-D**) Relative mRNA level of Notch target genes (*HES1*, *HEY1*, *CCND1*, and *APC*) (C) and Hippo target genes (*Axl*, *XIAP*, *Cyr61*, and *Myc*) (D). (**E**) PLA was performed in MGG6 GSCs to test close-proximity interactions between NR1D2 and TAZ, TEAD, Yap, RBPJ, and NICD1. (E’) Quantification of PLA signal intensity in (E) (n = 7, 7, 6, 6, 6, 7). (**F-G**) Co-IP assays to detect physical interactions between NR1D2-HA and endogenous TEAD, TAZ, NICD1, and RBPJ in U87 MG cells. Lysates from U87 MG cells with stably transfected NR1D2-HA were immunoprecipitated (IP) and probed with the indicated antibodies. (**H**) Cut&Tag q-PCR analysis of Axl, HEY1, and TCF7 in MGG6. MGG6 cells transfected with *HA-NR1D2* were used for HA enrichment quantification on promoter region (−500 to 0). (**I**) Schematic diagram of GSCs brain injection. (**J**) Survival curve of NSG mice bearing intracranial tumors from MGG6 GSCs transfected with *shNT, shNR1D2#1*, or *shNR1D2#2*, respectively. (**K**) *In vivo* bioluminescence imaging of NSG mice bearing tumors on day 7, 14, 17, 21, 23, 26, 30, 33, and 36 after MGG6 GSCs were injected into the mice brains. The injected MGG6 GSCs were transfected with *shNT*, *shNR1D2#1,* or *shNR1D2#2* respectively. (K’) Quantification of tumor size by *in vivo* luciferase assays. Statistical analysis by Unpaired *t* tests (A), two-tailed Student’s *t* tests (C, D, H), Two-way ANOVA analysis (B), or Log-rank (Mantel-Cox) test (J); Mean with SD. \**p* < 0.05, \*\**p* < 0.01, *** *p* < 0.001, **** *p* < 0.0001; Scale bar: 10 μm (E).

## Discussion

Hormone therapies are commonly employed to inhibit the proliferation of hormone-dependent cancers, such as breast and prostate cancers. Nonetheless, as time passes, cancer cells may develop mechanisms to survive, even in the absence of hormones or in the face of hormone blockers, resulting in resistance to hormone therapy.^1,7^ However, the specific mechanisms responsible for tumor progression and malignancy in hormone signaling-reduced tumors remain predominantly uninvestigated. Here we identified the primary hormone responsive gene *E75* as a crucial gene in promoting tumor malignant transformation in *Drosophila*. We found that inhibition of ecdysone signaling is a key characteristic of *Drosophila* malignant tumors, and inhibition of ecdysone signaling by ectopic expression of *E75* could facilitate the tumor malignancy of benign tumors by integrating two essential tumor-regulating pathways, namely the Hippo and Notch pathways (Fig. 7).

**Fig. 7.**
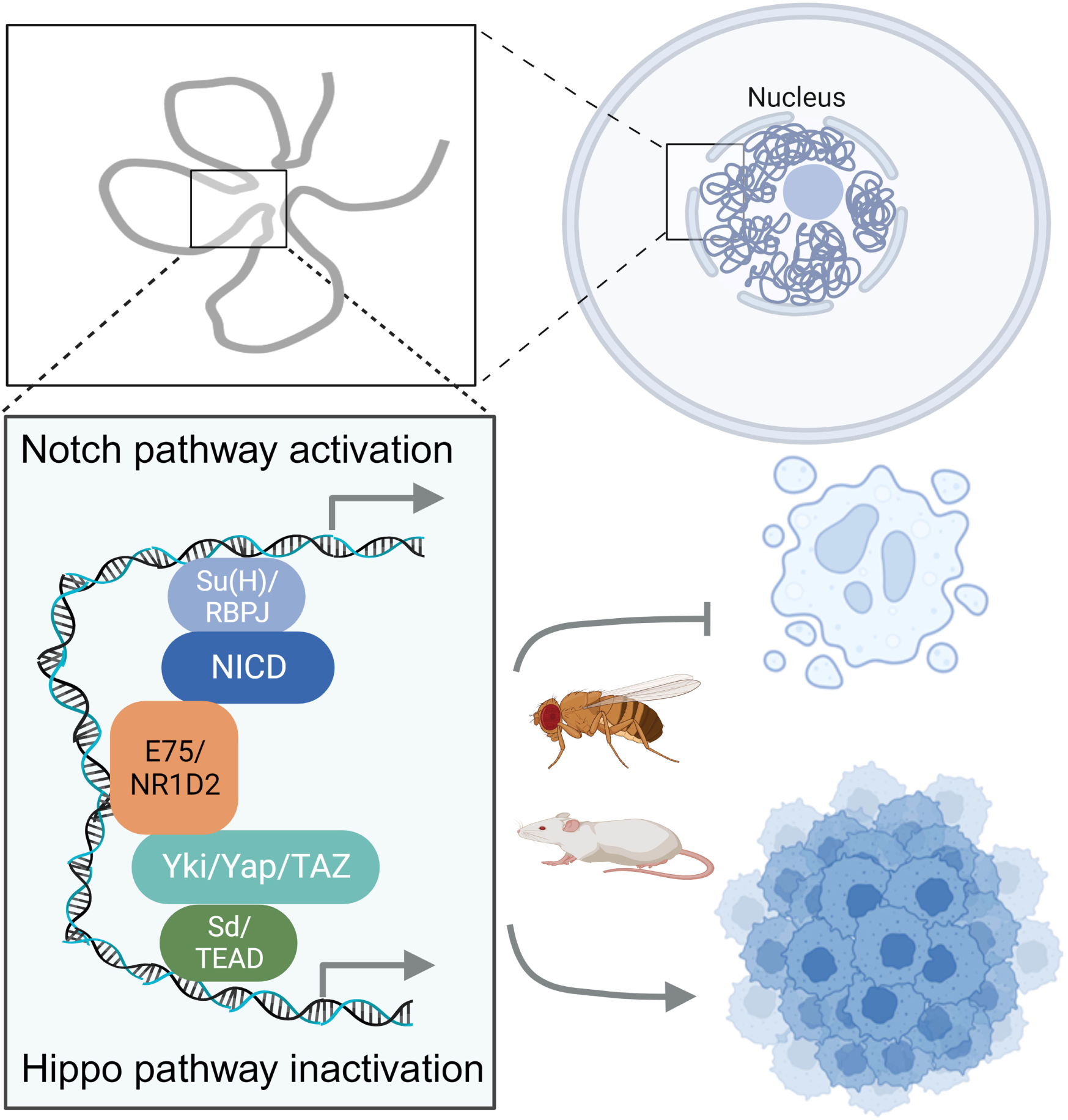
Model.

The nuclear receptor E75 has been implicated in the regulation of various biological processes, including axon degeneration and regrowth,^49^ circadian clock regulation,^50^ and cell migration.^24^ Recent studies have shown that ecdysone, released from the adult ovary following mating, exerts a remote effect on intestinal cell differentiation and proliferation through the downstream response gene *E75*.^22,23^ In this study, we reveal a novel growth and tumor-regulating role of E75 in *Drosophila* epithelium cells. Under physiological condition, depletion of *E75* induces apoptosis, while inhibition of ecdysone via *E75* overexpression promotes overgrowth. This is in line with a recent study showing that low titer ecdysone signaling activation promotes cell proliferation.^51^ Under pathological conditions, we found that ecdysone signaling inhibition induces tumor malignant transformation in diverse tumor models. Moreover, consistently with growth suppression role of ecdysone activation, we also show that hyperactivation of EcR effectively impedes *lgl^−/−^/Ras^V12^*-induced tumor malignancy.

The physical associations observed between E75 and the downstream transcription complexes of the Hippo and Notch pathways indicate that E75 potentially functions as an integrator, bridging these two signaling networks. We believe that this E75-mediated integration enables cells to generate coordinated responses that are essential for various physiological and pathological processes. Through the integration of Cut&Tag with RNA-seq analyses, we found that both NICD and Yki bind to the regions accessible to E75. Furthermore, E75 specifically binds and induces the expression of additional genes, including core transcription factor of the Toll signaling pathway. This suggests that, aside from its role in regulating the Hippo and Notch pathways, E75 may also serve as an integrator, coordinating the activity of multiple signaling pathways.

We have previously shown that expression of NR1D2 (also known as REV-ERBβ), the mammalian homolog of E75, was significantly upregulated in GBM cell lines.^45^. However, the specific pathways and biological functions in which NR1D2 is implicated remain poorly understood. Here we demonstrate that *NR1D2* expression is elevated in the GSCs, and the depletion of *NR1D2* inhibits GSC proliferation and improves the survival of tumor-bearing mice, indicating that NR1D2 plays an essential role in the progression of GBM *in vivo*. Moreover, we show that NR1D2, similar to E75, physically interacts with transcription factors of both Hippo and Notch pathways, regulating the expression of their target genes. This suggests an evolutionarily conserved role for E75/NR1D2 in tumorigenesis. It is noteworthy that pharmacological activation of *NR1D2* can induce cancer cell death, independent of NR1D2 function. As a result, treatment with NR1D2 agonists has been considered a promising anti-tumor strategy.^52–54^ However, as a result of this discrepancy, further examination is required to clarify the in vivo functions of NR1D2 by utilizing knock out mice in various tumor types, which could potentially provide insights that can advance the discovery of novel therapeutic strategies for specific cancers.

## Acknowledgements

We thank Jiong Chen, Oren Schuldiner, Tian Xu, Lei Xue, Duojia Pan, Suning Liu, Sheng Li, Zizhang Zhou, Hai Huang, Lei Zhang, Bloomington *Drosophila* Stock Center, Vienna *Drosophila* Resource Center, and Developmental Studies Hybridoma Bank for providing fly stocks and reagents; the Microscopy Core Facility and the High-Performance Computing Center of Westlake University for the facility support and technical assistance; Wenhan Liu for fly stock maintenance. This project was supported by National Natural Science Foundation of China (32170824, 32322027) to X.M. and (82073268) to Q.X., HRHI program (1011103360222B1) of Westlake Laboratory of Life Sciences and Biomedicine to X.M., Westlake Education Foundation to Q.X., “Pioneer” and “Leading Goose” R&D Program of Zhejiang (2024SSYS0034, 2024SSYS0036).

## Author Contributions

X.M. conceived and designed the study; X.M., X.W., Y.G., and Q.X. designed the experiments and analyzed the data; X.W. performed majority of fly related experiments with the help from S.S., W.X., M.W., and Y.W.; Y.G. performed bulk RNA-seq and CUT&Tag analyses with the help from Y.Z.; P.L. performed GBM-related experiments with the help from X.W., M.Y., and F.L.; X.M. and X.W. wrote the manuscript with input from Q.X., Y.G, and P.L.

## MATERIALS AND METHODS

### Key resource table

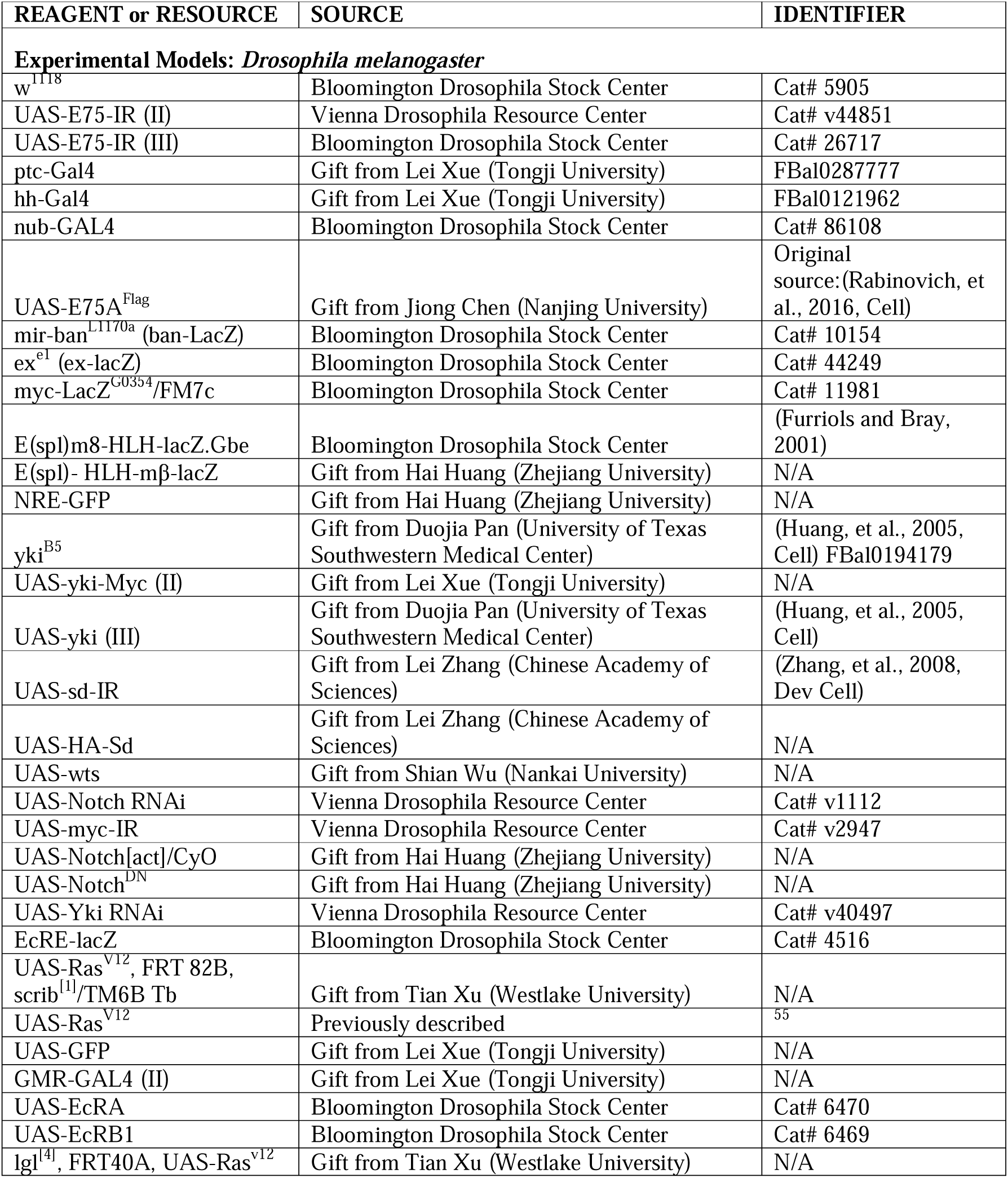
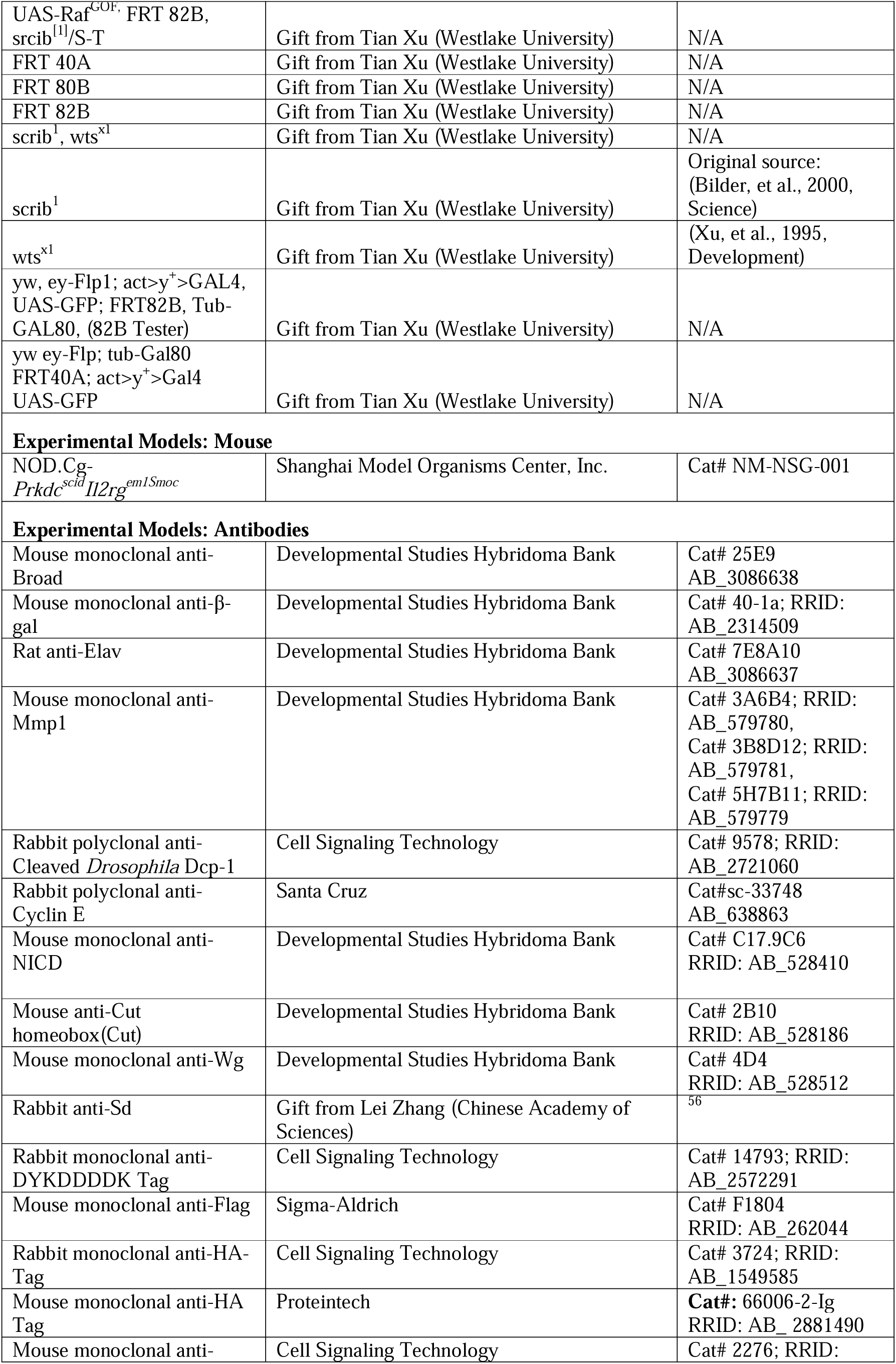
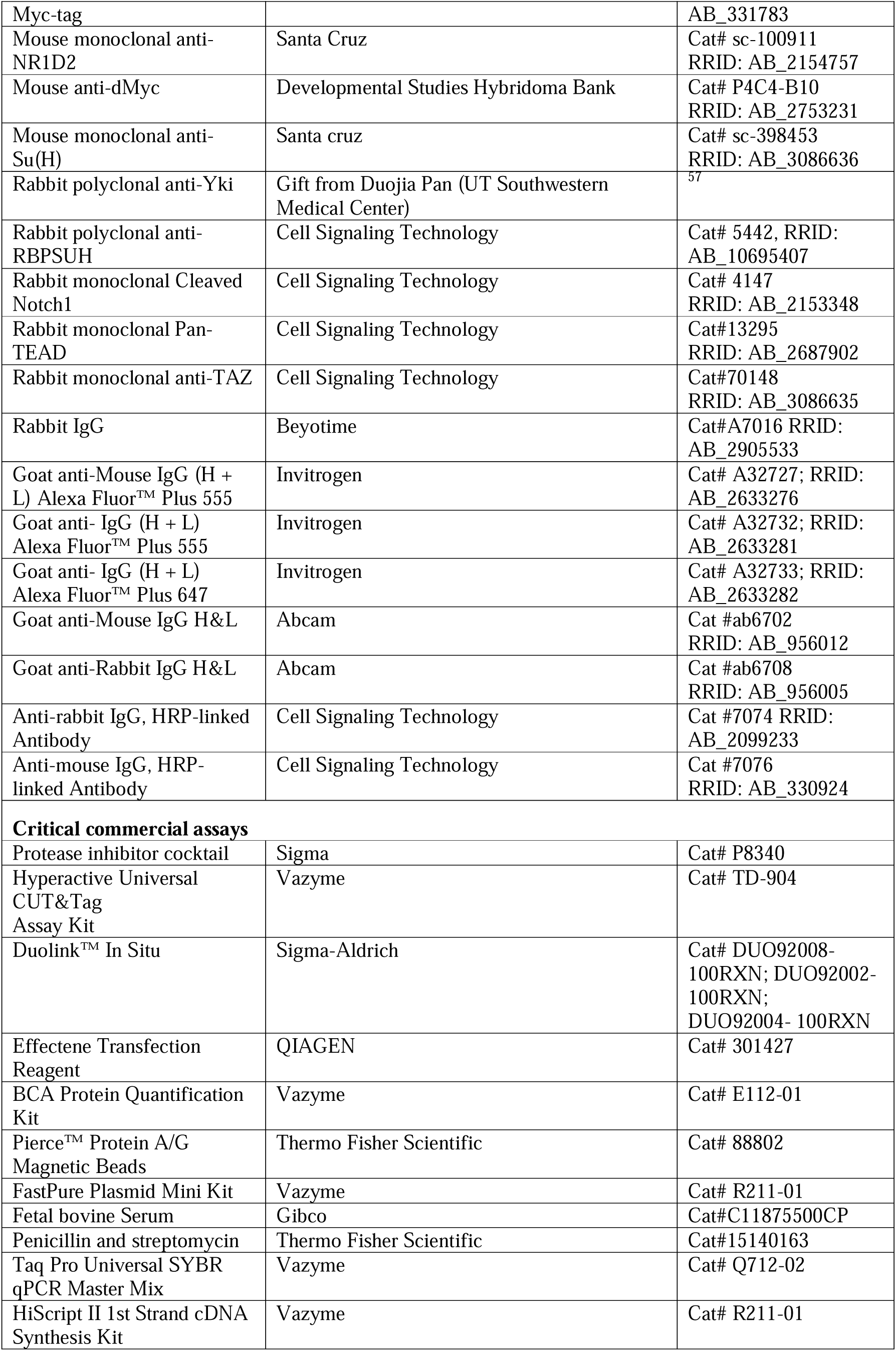
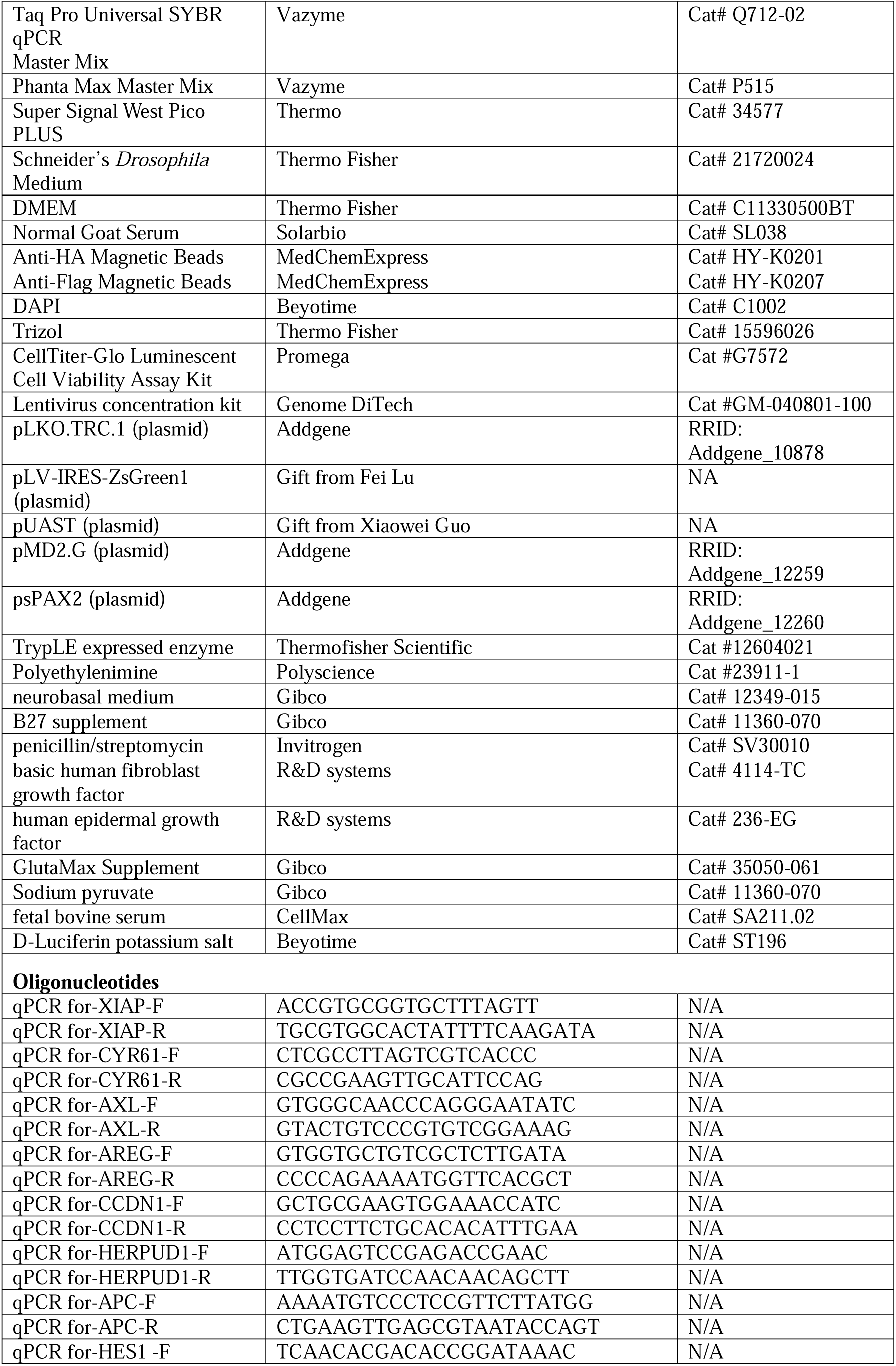
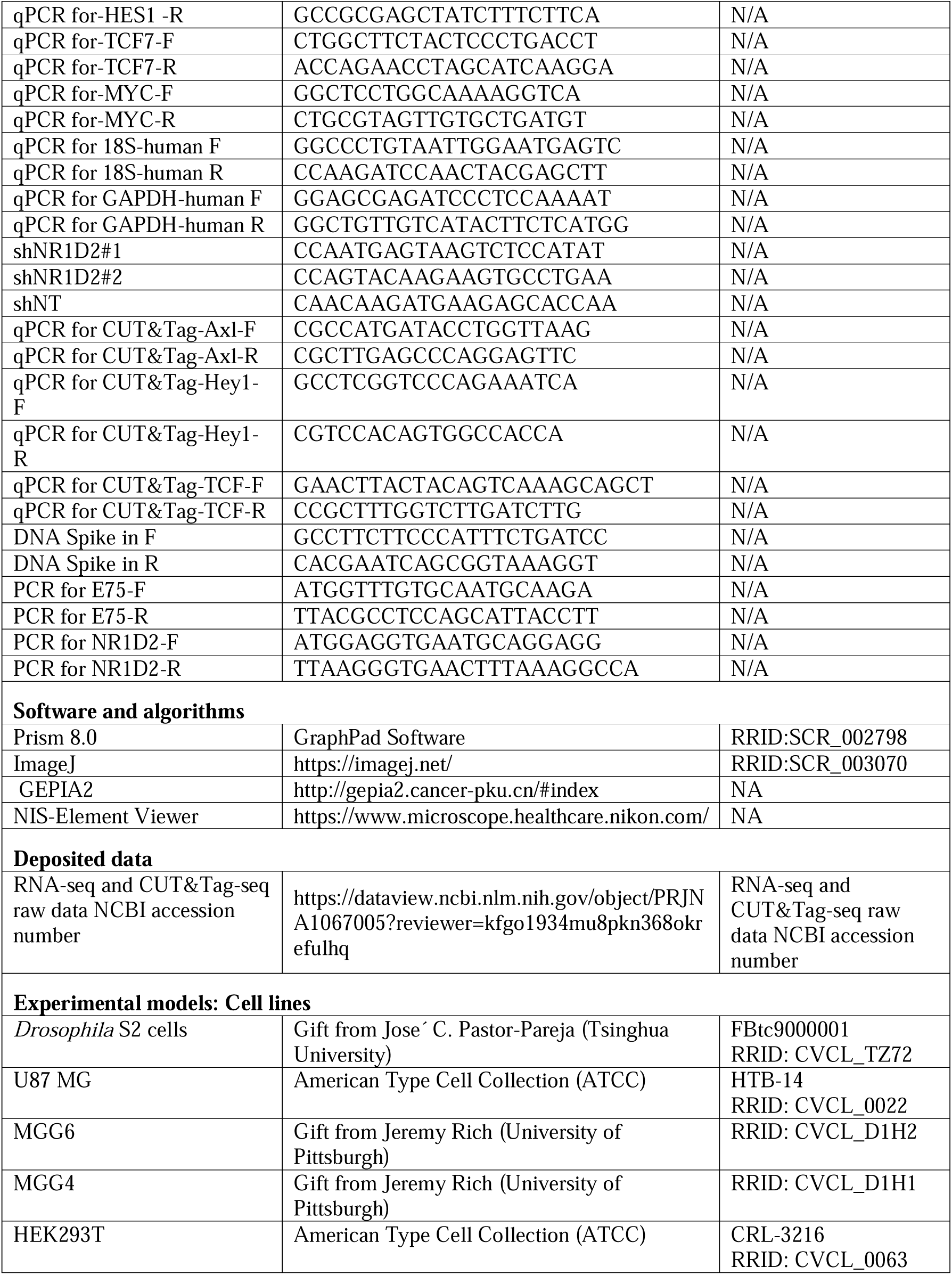

### STAR Methods

#### Fly husbandry and genetics

*Drosophila* stocks and crosses were maintained on standard food at a temperature of 25°C, unless otherwise specified. A comprehensive list of all Drosophila lines utilized can be found in the key resources table, and detailed genotypes for each figure panel are provided in Table S1. The standard food composition consisted of a cornmeal-yeast mixture, comprising 50 g of corn flour, 30 g of brown sugar, 24.5 g of dry yeast, 7.25 g of white sugar, 9 g of agar, 4.4 mL of propionic acid, 12.5 mL of ethanol, and 1.25 g of nipagin per liter.

#### Co-IP and western blot

Tumor samples or cells expressing indicated constructs with corresponding genotypes were collected and lysed in NP40 buffer with PMSF. Subsequently, the resulting cell lysate was combined with pre-washed magnetic beads, and the mixture was subjected to gentle rotation at room temperature for 30 minutes. This allowed for efficient binding of the target proteins to the magnetic beads. A magnetic separation rack was utilized, facilitating quick and easy isolation of the beads. The pre-cleared lysate, devoid of magnetic beads, was then carefully transferred to a clean tube, ensuring the retention of the protein content for subsequent analysis. Then, add primary antibody or HA-conjugated beads to 200 μl cell lysate, and incubate overnight at 4 °C. After that, pre-washed magnetic bead was added to the lysate and antibody solution (omit this step for antibody-conjugated beads), the bound protein can be finally separated through the magnetic beads. Finally, the coprecipitated protein was eluted for western blot analysis. The proteins were separated using SDS-PAGE, then transferred onto PVDF membranes. To prevent non-specific binding, the membranes were blotted with 5% skim milk powder in TBST for 60 minutes. Following this, the membranes were incubated overnight at 4 °C with primary antibodies. The following antibodies were used: HA (CST, 1:1000), Flag (Proteintech, 1:1000), Myc (CST, 1:1000), NICD (DSHB, 1:1000), Yki (1:1000), TEAD (CST, 1:500), RBPJ (CST, 1:500), TAZ (CST, 1:500), and NICD1(CST, 1:500). The membranes were washed three times with TBST to remove any unbound antibodies. Afterwards, the membranes were incubated with HRP-conjugated secondary antibodies to enable visualization of the target proteins. The following antibodies for western blotting were used: primary antibodies, rabbit-anti-FLAG (1:6,000), and secondary antibody anti-rabbit-HRP (1:8,000).

#### Duolink in situ proximity ligation assay (PLA)

Wing discs and cultured cells were harvested and subsequently fixed. Following fixation, the samples were blocked to prevent non-specific binding, and then subjected to a series of washes. The indicated primary antibodies form different origin (mouse or rabbit) were applied and allowed to incubate with the samples. Primary antibodies used *Drosophila* tissues include mouse anti-Flag (Sigma, 1:200) and rabbit anti-HA (CST, 1:200), rabbit anti-Flag (CST, 1:200) and mouse anti-Su(H) (Santa Cruz, 1:100), mouse anti-NICD (DSHB, 1:100), rabbit anti-Sd (from Lei Zhang, 1:100), rabbit anti-Yki (from Duojia Pan, 1:100). Primary antibodies used in mammalian cells include mouse anti-NR1D2 (Santa Cruz, 1:100), rabbit anti-TAZ, rabbit anti-pan TEAD (CST, 1:200), rabbit anti-Yap (CST, 1:200), rabbit anti-RBPJ (CST, 1:200) and rabbit anti-NICD1 (CST, 1:200). To enable detection, secondary antibodies conjugated to complementary PLA (Proximity Ligation Assay) probes were added. The Duolink method was employed, involving hybridization, ligation, amplification, and detection steps, following the guidelines provided by the manufacturer. This ensured accurate and reliable detection of protein-protein interactions or protein localization. For control samples, primary antibodies were omitted, and only secondary antibodies were added, which were subsequently detected using standard methods.

#### Cell proliferation assay

A total of 2500 cells were plated in each well of the 96-well plates. Subsequently, the Cell Titer-Glo Luminescent Cell Viability Assay Kit (Promega, Cat #G7572) was employed to assess cell viability at the indicated time points following the manufacturer’s instructions.

#### Mouse experiments

All mouse experiments were conducted in accordance with the relevant guidelines and under an animal protocol approved by the Institutional Animal Care and Use Committee of Westlake University. The intracranial transplantation of GSCs followed a previously described method ^58^. Briefly, GSC spheres were dissociated into single cells with TrypLE expressed enzyme (Thermofisher Scientific, Cat#12604021), and 10,000 cells were injected into the right cerebral cortex of NSG (NOD.Cg-Prkdc scid Il2rg tm1Wjl /SzJ) immunocompromised mice (Shanghai Model Organisms Center, Inc.) individually. As to in vivo tumor growth comparison, mice were narcotized and then imaged by bioluminescence imaging at indicated time. In parallel survival experiments, mice were monitored until the development of neurological signs or morbidity symptoms.

#### ShRNA Plasmids

The shRNA sequences were inserted into the pLKO.TRC.1 plasmid (Addgene, Cat#10878, RRID: Addgene_10878, Addgene).

#### Lentivirus production

HEK293T cells were cotransfected with a lentiviral expression vector, the envelope plasmid pMD2.G (Addgene, Cat#12259, RRID: Addgene_12259), and the packaging plasmid psPAX2 (Addgene, Cat#12260, RRID: Addgene_12260) using polyethylenimine (PEI) (Polyscience, Cat # 23966-1) according to the manufacturer’s instructions. After 48 hours of transfection, lentiviral particles were harvested and concentrated using the lentivirus concentration kit (Genomeditech, Cat# GM-040801-100).

#### Immunofluorescence and imaging

Third-instar larvae imaginal disc were dissected in cold PBS and fixed with 4% paraformaldehyde for 15 min at room temperature, then washed 3×5 min with PBS containing 0.1% Triton X-100 solution (PBST). Samples were blocked in 10% goat serum in PBST for 30 min after fixed and washed, and then incubated with primary antibody at 4°C overnight. Primary antibodies used include mouse Anti-beta Galactosidase (1:100, DSHB), mouse anti-Br (1:100, DSHB), rat anti-Elav (1:100, DSHB), mouse anti-Mmp-1 (1:100, DSHB), mouse anti-wg (1:100, DSHB), Cyclin E (1:100, Santa Cruz), rabbit anti-Dcp1 (CST, 1:100), mouse anti-dMyc (1:50, DSHB), mouse anti-Cut (1:100, DSHB), mouse anti-NICD (1:100, DSHB). After that, samples were washed 3×10 min with PBST, incubated with secondary antibody and DAPI at 1:200 in PBST for 2 hours. The images were performed with Nikon A1R confocal Microscope. Images were processed with NIS-Element Viewer and Image J software.

#### Total RNA extraction and Quantitative RT-PCR

In this study, a total 1 million GSCs or a total of 100 eye-antenna-disc tumors or a total of 150 wing imaginal discs were harvested at the late stage of third instar larvae for total RNA extraction of each biological replicate across different genotypes by using Trizol (Invitrogen, Carlsbad, CA) according to the manufacturer’s instructions. The RNA quality examination was conducted as follows: the purity of the sample was determined by NanoPhotometer^®^ (IMPLEN, CA, USA), while the concentration and integrity of RNA samples were detected by using Agilent 2100 RNA nano 6000 assay kit (Agilent Technologies, CA, USA). Subsequently, the extracted RNA was reverse transcribed into complementary DNA (cDNA) using a cDNA reverse transcription kit (Vazyme). Taq Pro Universal SYBR qPCR Master Mix (Vazyme) and the corresponding primers were utilized for quantitative polymerase chain reaction (qPCR), and the qPCR reactions were performed on a Jena Qtower384G Real-Time PCR System. 18S and GAPDH was used as an internal control in mammalian cell and Spike-in served as internal control for CUT&Tag q-PCR.

#### Library preparation for RNA sequencing

For each sample, a total amount of 1-3μg RNA was used as the input for library preparation by strictly following the standard protocol of VAHTS Universal V6 RNA-seq Library Prep Kit for Illumina^®^ (NR604-01/02). Briefly, mRNA was purified by using poly-T oligo-attached magnetic beads from total RNA. The short fragments of mRNA were obtained by adding the fragmentation buffer. After the first strand of cDNA was synthesized by using random hexamer primer and RNase H, the second strand synthesis was performed subsequently by using buffer, dNTPs, DNA polymerase I, and RNase H. And then, the double-stranded cDNA was purified by using QiaQuick PCR kit or AMPure P beads. The purified products of each sample were repaired at the end, added tail, and connected to the sequencing connector, then the appropriate fragment size was selected, and the final cDNA library was obtained by PCR amplification for further sequencing performed on Illumina NovaSeq 6000 platform with NovaSeq 6000 S4 Reagent kit V1.5.

#### RNA-seq data processing and analysis

Both wing disc and eye-antennal disc transcriptomic data were processed and analyzed as follows: Data quality control and reads statistics were performed by using FastQC (v0.11.8) software (https://github.com/s-andrews/FastQC/releases/tag/v0.11.8). Low quality reads were removed, and the maintained high-quality reads (Q30 > 90%) were mapped to the Ensemble ^59^ *Drosophila melanogaster* reference genome (Drosophila_melanogaster.BDGP6.32.108) by using Hisat2 ^60^. HTSeq ^61^ was applied for gene feature counting with default settings. Differential gene expression identification and functional annotation were conducted in R (v4.2.0). Genes meeting |Fold-Change| > 1.5 and false discovery rate (FDR) < 0.05 were identified as differentially expressed genes (DEGs) by using edgeR ^62^ with “RLE” method. The gene id transformation was performed by using biomaRt ^63^. DEGs were annotated against terms in Kyoto Encyclopedia of Genes and Genomes (KEGG) database ^64,65^ and Gene Ontology (GO) consortium ^66^ and by using clusterProfiler package ^67^. Data visualization was relied on pheatmap (v1.0.12), RColorBrewer (v1.1-3), and ggplot2 (v3.4.4) packages. Gene set enrichment analysis (GSEA) ^68,69^ was performed locally with gene list of each pathway from FlyBase (https://flybase.org) ^70^.

#### Cleavage under targets and tagmentation (CUT&Tag)

All CUT&Tag experiments were conducted by strictly following the manual instructions of Hyperactive Universal CUT&Tag Assay Kit for Illumina Pro (Vazyme, TD904). Briefly, approximately 1 x 10^5^ cells were collected from wing imaginal disc or eye-antennal disc tumors for each sample group (at least 3 biological replicates). These cells were incubated and mixed (2-3 times) with Concanavalin A beads Pro at room temperature for 10 min. Then, the liquid was removed, and the specific primary antibodies were added with ice-cold 50μl Antibody Buffer. The primary antibodies used include Myc-tag (CST, 1:100), NICD(DSHB, 1:50), Flag-tag (Sigma, 1:50) in *Drosophila* tissues; Rabbit IgG (Beyotime, 1:100) or HA-tag (CST, 1:100) for Cut&Tag q-PCR in mammalian cells. After incubating overnight at 4L, the liquid was removed, and the secondary antibody diluted by Dig-Wash Buffer (1:100) was added for incubation at room temperature for 1 h. For each sample, 2μl Hyperactive pA/G-Transposon Pro mixed with 98μl Dig-300 Buffer were added and incubated at room temperature for 1 h. After gently washing with Dig-300 Buffer, the fragmentation of each sample was conducted by incubating the mixture of 40μl Dig-300 Buffer, 10μl 5 × TTBL, and the products obtained in the previous step at 37 for 1h. 1pg spike-in was added for internal control before DNA extraction. Then, DNA extraction was performed by incubating the products with DNA Extract Beads Pro diluting in 50μl 2 × B&W Buffer at room temperature for 20 min. The i7 and i5 Indexed Primer were combined for PCR amplification with recommended cycle number (9~11). Finally, only the CUT&Tag libraries passing fragment analyzer quality control (Fragment Analyzer-12/96) were then sequenced on Illumina NovaSeq 6000.

#### CUT&Tag-Seq data processing and analysis

The analytic procedures of CUT&Tag-seq data were referred to recommended pipelines.^71^ The raw sequencing data were processed by using FastQC (v0.11.8) software (https://github.com/s-andrews/FastQC/releases/tag/v0.11.8) for quality control. Adapters were removed and reads were trimmed by using Cutadapt (v1.18).^72^ The clean reads were first mapped to the Ensemble ^59^ *Escherichia coli* (E. coli) reference genome (GCF_000005845.2_ASM584v2_genomic) by using Bowtie2 (v2.4.2),^73^ the unrecognized reads were subsequently mapped to *Drosophila melanogaster* reference genome (Drosophila_melanogaster.BDGP6.32.108) with the parameters “--end-to-end --very-sensitive --no-mixed --no-discordant --phred33 -I 0 -X 1000 --no-unal”. After using SAMtools (v1.11)^74^ to transform SAM files into BAM files, Picard (v2.25.1) (https://broadinstitute.github.io/picard/) was applied to remove duplicates, and the reads were selectively maintained when meeting “mapping_quality >= 20”. The final BAM files of same genotype among the replicates were merged into one for peak calling, which is performed by using Model-based Analysis of ChIP-seq (MACS2, v2.2.6)^75^ with settings “-g dm -f BAMPE -q 0.05 --keep-dup all”. Peaks were annotated by using ChIPseeker^76^ package in R (v4.2.0), and the peak distribution was visualized by using *plotAnnoBar* and *plotDistToTSS* functions. The final BAM files were transformed into bigwig files for signal enrichment visualization by using the *bamCoverage* function of BEDTools (v2.30.0)^77^ with settings “--binSize 20 --normalizeUsing BPM”. The average normalized signal values of peaks in *drosophila* genome region were calculated and visualized by using DeepTools (v3.5.1),^78^ in which regions within 3kb distance relative to the transcriptional start sites (TSS) were included and gene body region were scaled into 5kb length. Integrative Genomics Viewer (IGV)^79^ was applied to visualize the peaks on specific genome region of interested genes. The binding motifs were identified by using the *findMotifsGenome* or *findMotifs* functions of Homer (v4.11)^80^ regarding to conditions. The motif scanning algorithm, Find Individual Motif Occurences (FIMO, v5.5.5),^81^ was applied in a set of sequences to determine all the positions where transcription factor motifs match (*p* < 0.0001) via The MEME Suite ^82^.

### Public data analysis

The binding regions of Yki and Su(H) on *Drosophila melanogaster* genome were annotated by using public datasets GSE38594^83^ and GSE41429,^84^ respectively. The DEG list between siNR1D2 and siControl LN-18 cells was retrieved from the Supplementary Table S1 and S2 of its publication,^85^ and the over-representation analyses (ORA) were conducted using clusterProfiler package.^67^ The analysis of *NR1D2* relative expression level between GSC and NSC was conducted by using the processed data (CPM) from GSE54791^86^ for the Mann-Whitney test. The correlation analysis of *NR1D2* expression with other genes of interest were performed by using the GEPIA2 web server.^87^

### Data and code availability

All sequencing data of this study is deposited in the Sequence Read Archive (SRA) with the accession number PRJNA1067005 (https://dataview.ncbi.nlm.nih.gov/object/PRJNA1067005?reviewer=kfgo1934mu8pkn368okrefulhq). All code for data cleaning and analysis associated with the current study is available upon request.

### Quantification and statistical analysis

In Figure 1 F, G and Figure S1C, the region of interest (ROI) in each clone and the adjacent wild-type cell was subjected to cycling and measurements using ImageJ. The size of the tumors or ROI were quantified by measuring the area of the GFP-positive region in each sample. All statistical analyses were performed with GraphPad Prism 8.0. software. Data represent mean values ±SD. Statistical significance was assessed using appropriate methods depending on the experimental design. For comparisons between two groups, an unpaired two-tailed Student’s t-test was performed. For experiments involving three or more groups, Ordinary one-way ANOVA or two-way ANOVA analysis was conducted. Additionally, in survival analysis, the Log-rank (Mantel-Cox) test was employed. The specific statistical tests used for each analysis are indicated in the corresponding figures. *p* value < 0.05 was considered significant, \**p* < 0.05, \*\**p* < 0.01, \*\*\**p* < 0.001, \*\*\*\**p* < 0.0001.

