## Supplemental data for "Nuclear receptor E75/NR1D2 drives tumor malignant transformation by integrating Hippo and Notch pathways"

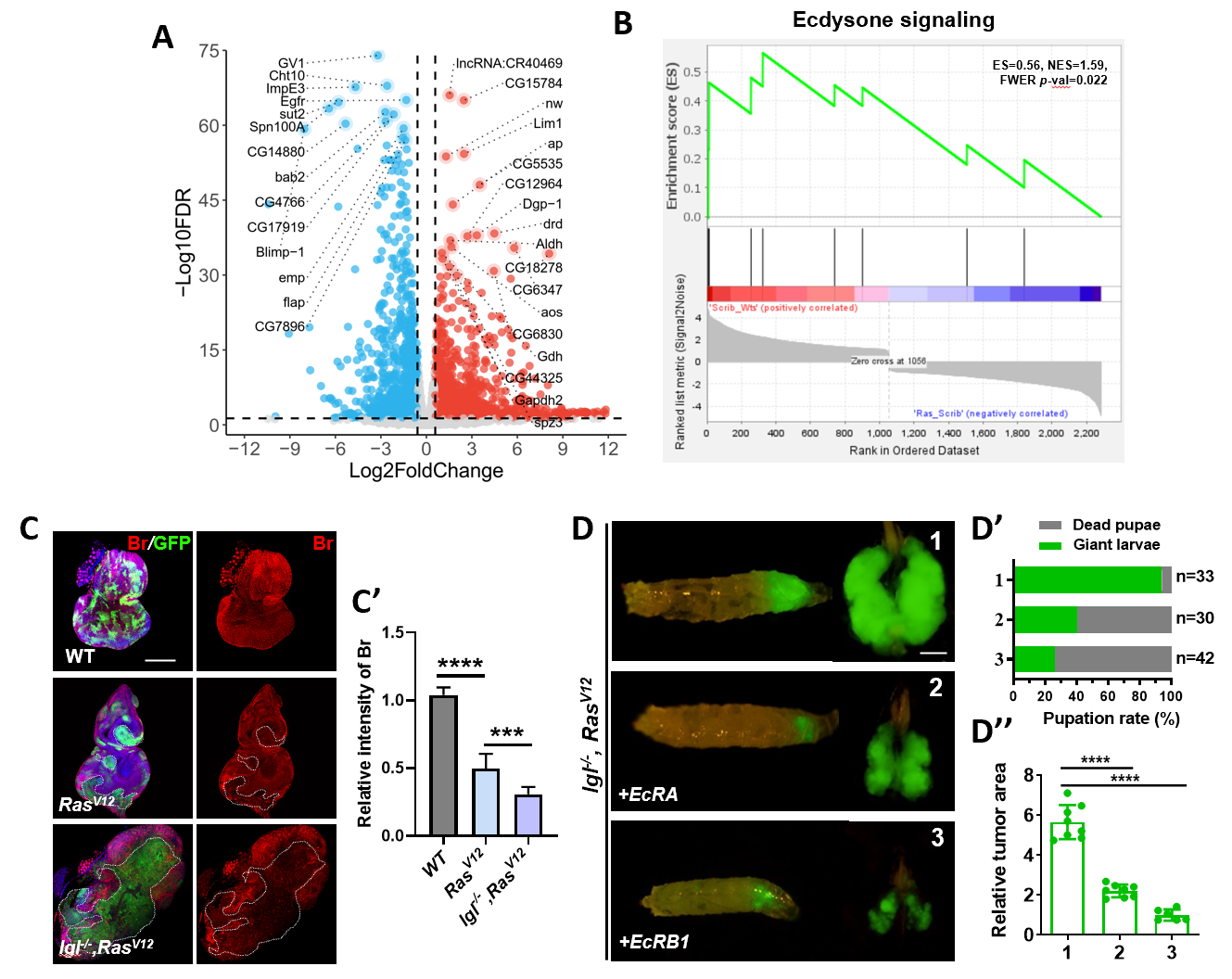

**Fig. S1 Ecdysone signaling is inhibited in malignant tumor, related to Figure 1.**

(A) Volcano plot of differentially expressed genes (DEGs) in *scrib^-/-^,wts^-/-^* and *scrib^-/-^,Ras^v12^* tumors.

(B) GSEA enrichment of ecdysone-related genes in *scrib^-/-^,wts^-/-^* and *scrib^-/-^,Ras^v12^* tumors.

(C) Confocal images of eye-antennal discs bearing *ey-Flp-*MARCM-induced mosaics of each genotype stained with Broad (Br) antibody. (G’) Quantification of relative Br intensity of GFP positive mosaics clones (n = 6, 7, 8).

(D) Representative images of tumor-bearing larvae and *ey-Flp-*MARCM-induced tumors with indicated genotype. Quantification of pupation rate of tumor-bearing larvae (D’) and relative tumor size of GFP positive mosaics clones (D’’).

Statistical analysis by Ordinary one-way ANOVA test; mean ± SD. ***p < 0.001, ****p < 0.0001. Scale bars:100 μm (C), 200 μm (D).

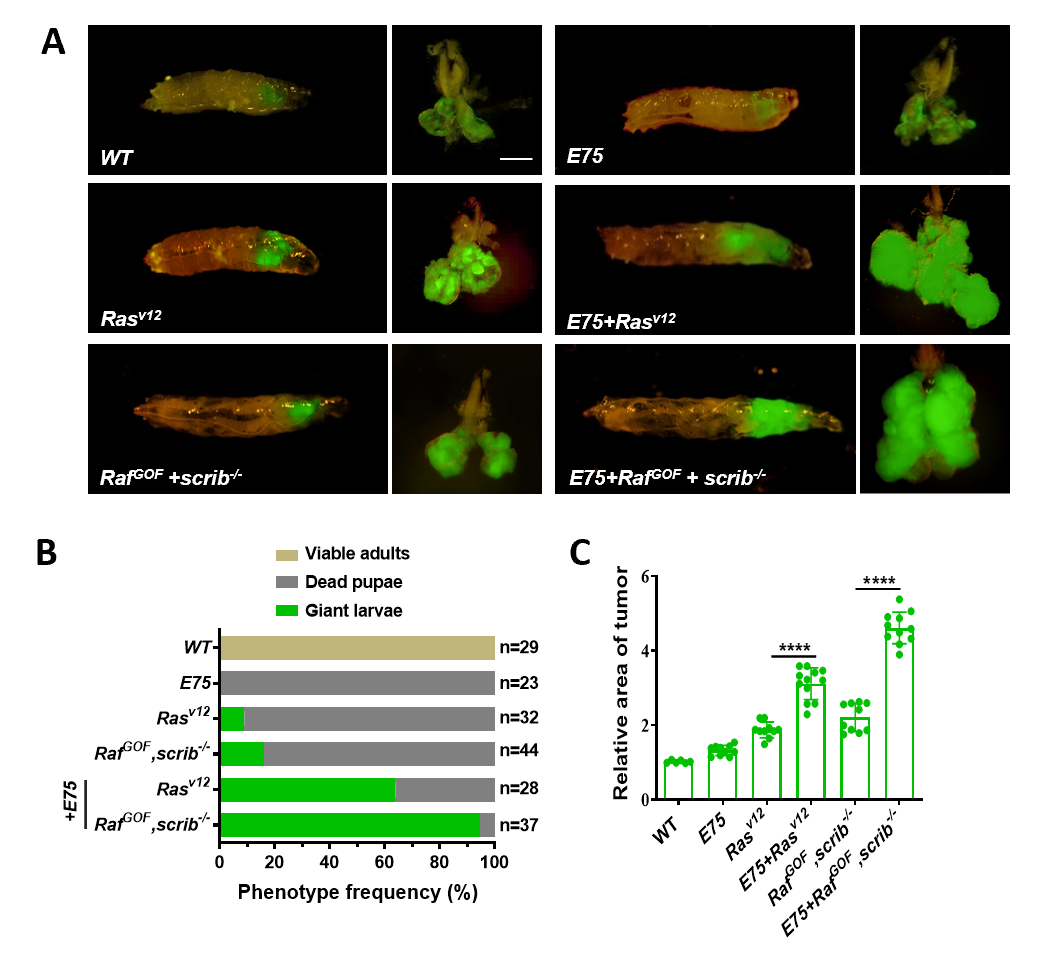

**Fig. S2 *E75* overexpression induces tumor malignant transformation, related to Figure 2.**

(A) Dorsal views of ey-Flp-MARCM-induced GFP-positive tumor-bearing larvae and the corresponding eye disc or tumor (right).
(B) Quantification of larvae pupation rate in A.
(C) Quantification of relative tumor size in A (n = 10, 10, 12, 10, 11).

Statistical analysis by Ordinary one-way ANOVA test. mean ± SD. ****p < 0.0001. Scale bars: 200 μm.

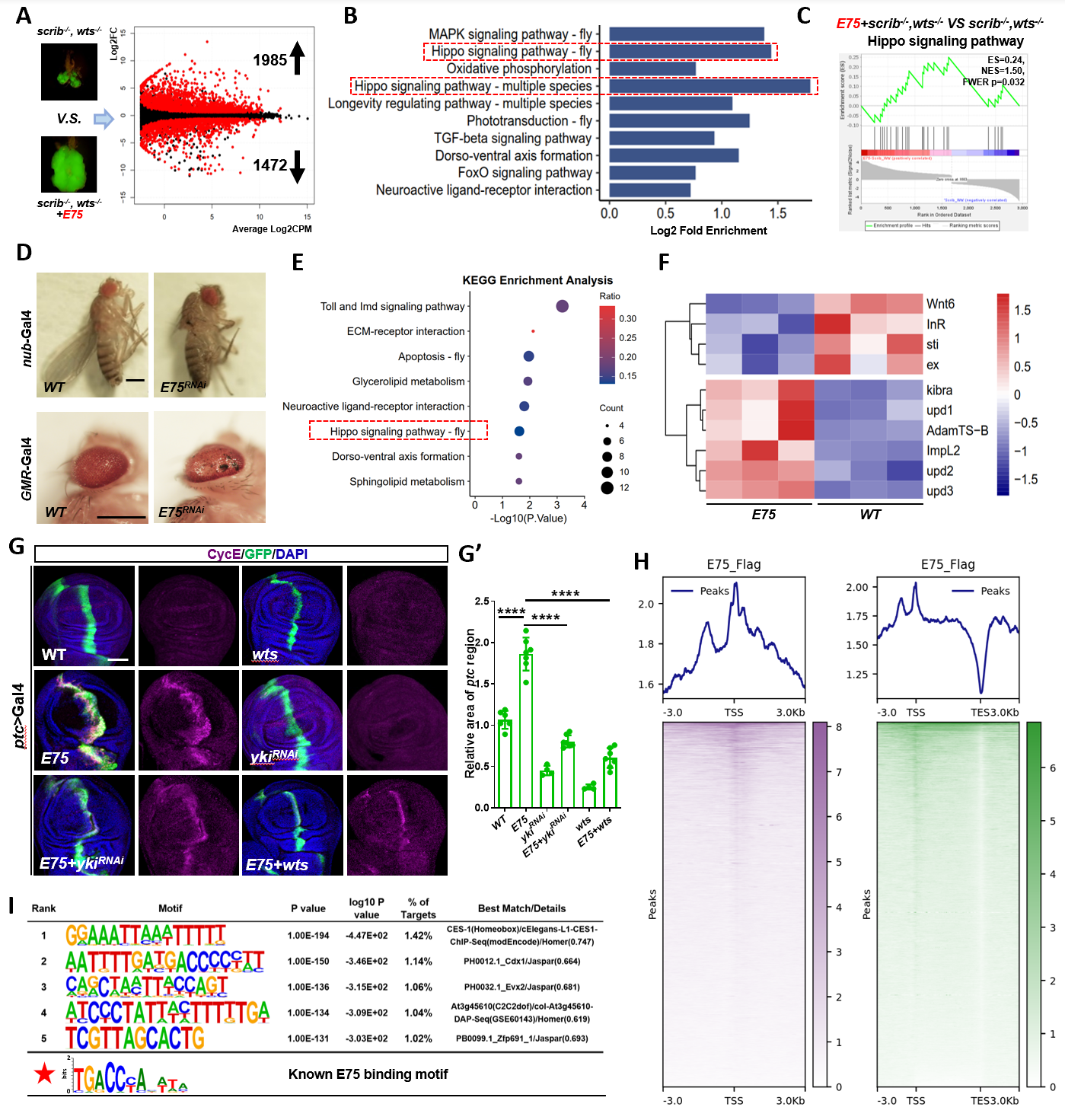

**Fig. S3 *E75* inactivates the Hippo pathway to induce tumor malignancy, related to Figure 3.**

(A) MA (Minus-versus-Add) plot of *scrib^-/-^, wts^-/-^* and *E75,scrib^-/-^, wts^-/-^* tumors.

(B) Enrichment analysis of signaling pathway in *scrib^-/-^,wts^-/-^* and *E75,scrib^-/-^, wts^-/-^* tumors.

(C) GSEA enrichment of Hippo signaling-related genes in *scrib^-/-^, wts^-/-^* and *E75, scrib^-/-^, wts^-/-^* tumors.

(**D**) Light micrographs of the adult wings (top) and eyes (bottom) bearing the indicated genotypes are shown.

(**E**) KEGG enrichment of signaling pathways in *scrib^-/-^, wts^-/-^* and *E75, scrib^-/-^, wts^-/-^* tumors.

(**F**) Heatmap profiles of Hippo signaling-related genes in wild-type (WT) and *E75* overexpression wing discs.

(G) Confocal image of cyclin E (CycE) antibody staining of wing discs bearing the indicated genotypes. (G’) Quantification of the relative size of *ptc* region of G.

(H) Binding profiles and heatmaps of E75 Cut&Tag signals are displayed within a region spanning ± 3 kb around all canonical transcription start sites (TSS) (left), -3 kb around all TSS and +3 kb around all canonical transcription end sites (TES) (right).

(**I**) HOMER motif analysis of E75 binding regions.

Statistical analysis by Ordinary one-way ANOVA test. mean ± SD. ****p < 0.0001. Scale bars: 100 μm (G), 200 μm (D).

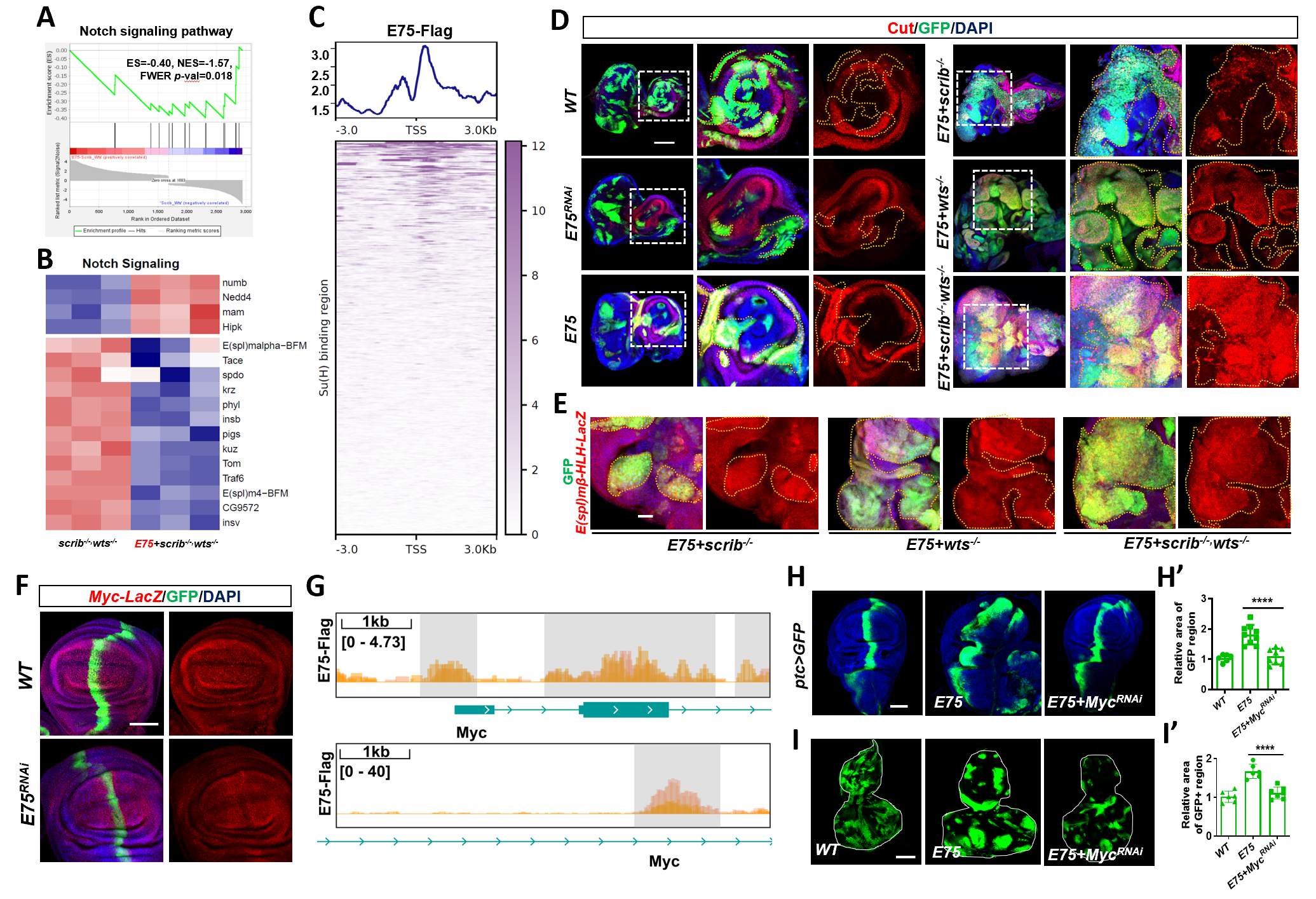

**Fig. S4 E75 activates the Notch pathway to promote tumor malignancy, related to Figure 4.**

(A, B) GSEA enrichment (A) and heatmap profiles (B) of Notch signaling-related genes in *scrib^-/-^, wts^-/-^* and *E75, scrib^-/-^, wts^-/-^* tumors.

(C) Line plots of the average Cut&Tag signal of E75 (top panel) and the heatmaps of the Cut&Tag signals of Su(H) in *Drosophila* (bottom panel). Cut&Tag signals are displayed within a region spanning ± 3 kb around all canonical transcription start sites (TSS) genome-wide.

(D, E) Representative confocal images of Cut (D) and *E(spl)mβ-HLH*-*lacZ* (E) staining in *ey-Flp-*MARCM-induced tumors and clones with indicated genotype.

(F) Representative confocal images of *Myc-lacZ* staining in wing discs of wild-type and *E75* knockdown.

(**G**) Browser shots of *E75* CUT&Tag signal at the regulatory region of *Myc*.

(**H, I**) Representative confocal images of wing (H) and eye discs (I) with indicated genotypes. (H’, I’) Quantification of relative GFP region of H and I (H’, n = 6, 9, 8; I’, n = 6, 6, 7).

Statistical analysis by Ordinary one-way ANOVA test. mean ± SD. ****p < 0.0001. Scale bars: 100 μm.

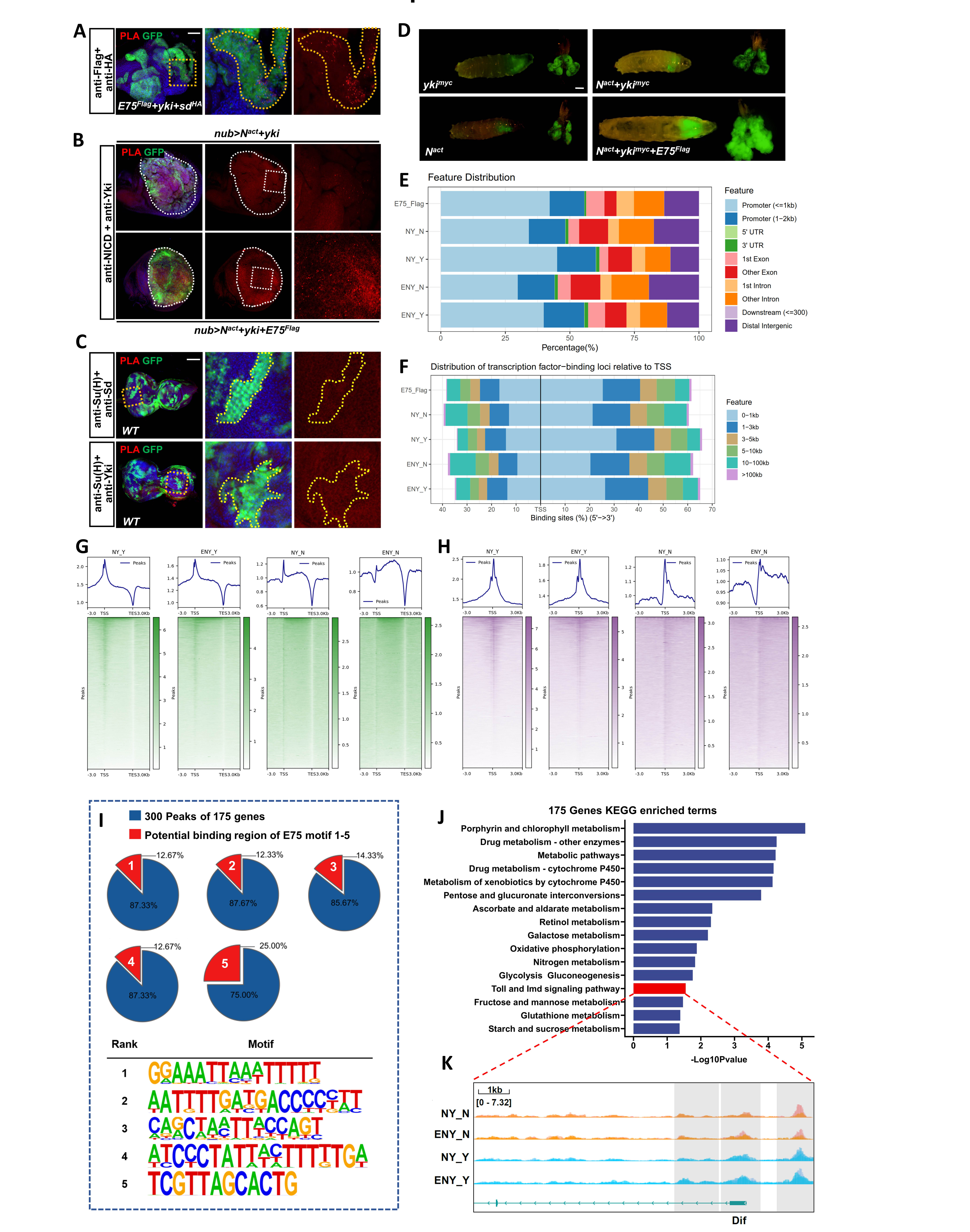

**Fig. S5 E75 integrates the Hippo and Notch pathways at the transcription factor level, related to Figure 5.**

(A) PLA was performed on eye discs bearing *ey-Flp-*MARCM-induced *E75^Flag^*, *Yki*, and *Sd^HA^* co-expressed clones to test close-proximity interactions between E75 and Sd.

(B) PLA was performed on wing discs with *Yki* and *Sd^HA^* co-expression under the control of *nub* promoter, with or without *E75^Flag^* expression, to test close-proximity interactions between NICD and Yki.

(C) PLA was performed on eye discs bearing *ey-Flp-*MARCM-induced WT clones to test close-proximity interactions between Su(H) and Sd, as well as Su(H) and Yki.

(D) Dorsal views of ey-Flp-MARCM-induced GFP-positive tumor-bearing larvae and the corresponding eye disc or tumor (right).

(E) Feature distribution of genomic annotations of Cut&Tag in E75 peaks (E75_Flag), NICD peaks of *N^act^* and *Yki* overexpression (NY_N), Yki peaks of *N^act^* and *Yki* overexpression (NY_Y).

(F) Distribution of binding loci relative to TSS of peaks mentioned in E.

(G) Binding profiles and heatmaps of NICD and Yki in *N^act^,Yki* and *E75, N^act^,Yki* tumors. Cut&Tag signals are displayed within a region spanning -3 kb around all canonical TSS and +3 kb around all canonical transcription end sites (TES).

(H) Binding profiles and heatmaps of NICD and Yki in *N^act^,Yki* and *E75, N^act^,Yki* tumors. Cut&Tag signals are displayed within a region spanning -3 kb around all canonical TSS and +3 kb around all canonical TSS.

(**I**) Percentage of five E75 motifs in 300 peaks of 175 genes mentioned in Figure 5J.

(**J**) KEGG enriched terms of 175 genes mentioned in Figure 5J.

(**K**) Browser shots of NICD and Yki CUT&Tag signal at the regulatory region of *Dif* in NY and ENY.

Scale bars: 100 μm (A, B, C), 200 μm (D).

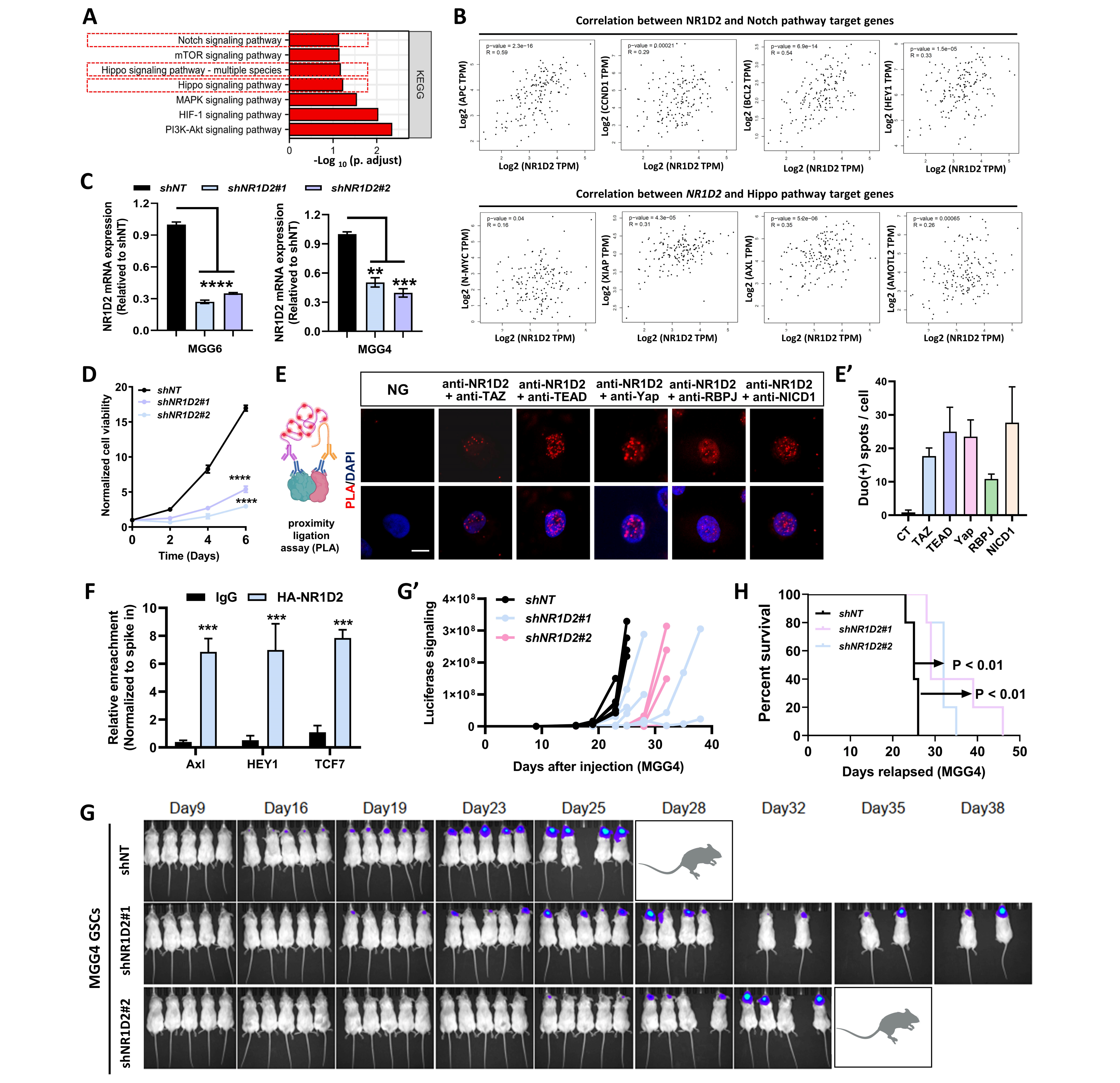

**Fig. S6 Silencing NR1D2 suppressed glioblastoma stem cell-driven tumor growth, related to Figure 6.**

(A) KEGG analysis of *NR1D2*-depleted U87 MG cell.

(B) Pair-wise gene correlation analysis of *NR1D2* and target genes of Notch pathway (top panels) and Hippo pathway (bottom panels) in GBM by GEPIA2.

(C) Relative mRNA expression of *NR1D2* in GSCs and *NR1D2*-depleted cells.

(D) Relative cell viability of MGG4 of shNT and *shNR1D2*.

(E) PLA working model (left). PLA analysis in U87 MG cells to test close-proximity interactions between NR1D2 and TAZ, TEAD, Yap, RBPJ, and NICD1 (right). (E’) Quantification of PLA signal intensity in E (n = 7, 6, 9, 6, 6, 6).

(F) Cut&Tag q-PCR analysis of *Axl*, *HEY1*, and *TCF7* in U87 MG. U87 cells transfected with *HA-NR1D2* were used for HA enrichment quantification on promoter region (−500 to 0).

(G) *In vivo* bioluminescence imaging of NSG mice bearing tumors on day 9,16, 19, 23, 25, 28, 32, 35, and 38 after MGG4 GSCs were injected into the mice brains. The injected MGG4 GSCs were transfected with *shNT*, *shNR1D2#1*, or *shNR1D2#2*, respectively. (G’) Quantification of tumor size by in vivo luciferase assays.

(H) Survival curve of NSG mice bearing intracranial tumors from MGG4 GSCs transfected with *shNT*, *shNR1D2#1*, or *shNR1D2#2*, respectively.

Statistical analysis by two-tailed Student’s t tests (C, F), two-way ANOVA (D), or Log-rank (Mantel-Cox) test (H); ***p* < 0.01, ****p* < 0.001, *****p* < 0.0001; Scale bar: 10 μm (E).
